# Plasticity in a bacterial global regulatory switch that drives a shift in antibiotic resistance and virulence

**DOI:** 10.1101/2025.06.09.658697

**Authors:** Annapaula Correia, Emma J. Manners, Benjamin A. Evans, Jacob G. Malone, Justin O’Grady, Eleni Mavrogiorgou, Sergio Arminu, Manuela Usai, Dixita Naik, Stuart McMillan, Gemma L. Kay, Claire Hill, Padhmanand Sudhakar, Emma Meader, Katarzyna Schmidt, Tamas Korcsmaros, Andrew P. Desbois, Lisa Crossman, John Wain, Gemma C. Langridge

## Abstract

Antibiotic resistance and expression of virulence factors can impact the outcome of infection by *Pseudomonas aeruginosa*. Pathogenesis is often modelled using the *P. aeruginosa* PAO1 reference strain but laboratory lineages vary in the sequence and activity of MexT, a global regulator impacting virulence, biofilm formation and ciprofloxacin resistance. We defined the impact of active versus inactive MexT in PAO1 and observed global transcriptomic changes affecting the expression of 900 genes. Phenotyping revealed altered metabolism, antibiotic resistance and virulence - resulting in striking variation across a ‘single’ model organism. We propose that antibiotic-resistance, introduced during work with the original PAO1 strain, has caused plasticity in *mexT* that accounts for variation across lineages. To test this, we introduced antibiotic resistance into clinical *P. aeruginosa* isolates and observed downstream mutations in *mexT* when selective pressure was removed, supporting the proposed evolutionary pathway. Overall, we have demonstrated the transcriptomic basis of MexT as a phenotypic switch in PAO1 and implicated antibiotic resistance as a cause of downstream changes in *mexT* in *P. aeruginosa*. Furthermore, MexS/MexT-regulated efflux is implicated in the antibiotic stress response and virulence, helping identify the mechanisms for rapid phenotypic switching through *mexT* and confirming that PAO1 is a very different organism compared to most isolates. Improved understanding of the regulatory changes linked to antibiotic resistance is particularly relevant to *P. aeruginosa* where cycles of antibiotic treatment are common.

## Introduction

Bacterial survival is dependent on rapid adaptation to the ever-changing environments bacteria are exposed to. Rapid changes in gene regulation can result in phenotypic plasticity enabling bacteria to cope with environmental changes, whereas in other cases genetic changes are selected for that result in the new phenotype. Understanding how bacteria adapt to changes in the environment, such as exposure to antibiotics, is crucial for drug development and understanding how to best utilise drugs already available. Much is known about the properties of specific antibiotic resistance mechanisms, but less is known about the evolutionary trajectories and epistatic interactions that cause specific genotypes of antibiotic-sensitive naive strains to evolve into multidrug-resistant superbugs.

*Pseudomonas aeruginosa* is a WHO priority pathogen that is frequently multidrug-resistant (Prioritization of pathogens to guide discovery, research and development of new antibiotics for drug-resistant bacterial infections, including tuberculosis, 2017). Its adaptability means it can survive in contaminated soil and water, man-made water systems, and the debilitated human tissues of diabetic and cystic fibrosis patients (Crone *et al*, 2020). Innate and acquired antimicrobial resistance (AMR) in *P. aeruginosa* greatly increases the ability to evade antibiotics and infect the host (Langendonk, Neill and Fothergill, 2021).

The best described mechanisms of AMR in *P. aeruginosa* are enhanced extrusion systems (multidrug efflux, Mex) combined with reduced antibiotic ingress (Schweizer, 2003; Livermore, 2002). The 12 Resistance-Nodulation-Division (RND) systems are the most important of the recognised efflux pumps; some, such as MexAB-OprM, encode intrinsic resistance to toxic substances and are expressed constitutively (Li, Nikaido and Poole, 1995; Piddock, 2006). Other RND systems however, such as MexEF-OprN, are inducible and represent an adaptive response. MexEF-OprN contributes to trimethoprim, chloramphenicol and fluoroquinolone resistance through efflux (Maseda, Yoneyama and Nakae, 2000) and is counter-regulated with the outer membrane porin OprD which is permeable to imipenem and meropenem. This means that multi-drug resistance (MDR) can be associated with efflux and permeability barriers through a single regulatory event (Ochs *et al*., 1999). Importantly, the role of MexEF-OprN goes far beyond MDR as it is involved in amino acid/peptide uptake through OprD, export of natural toxins (Juarez *et al*., 2017; Fetar *et al*., 2011; Fargier *et al*., 2012) and quorum sensing (Köhler *et al*., 2001; Lamarche and Déziel, 2011; Dreier and Ruggerone, 2015) thus playing a role in host adaptation and bacterial cell-to-cell communication.

Understanding gene regulation is key to understanding biological success. The balanced control of MexEF-OprN in *P. aeruginosa,* through regulation by MexT and MexS (and potentially other proteins), exemplifies the complexity that has evolved in prokaryotic systems (Figure 1). Under standard laboratory conditions (i.e. antibiotic-free, rich media) MexT, a transcriptional activator for MexEF-OprN, is itself suppressed by MexS activity (Juarez *et al*., 2017). MexS is an oxidoreductase that is thought to detoxify electrophilic metabolites and maintain the cellular redox balance. This keeps MexT in an inactive form and efflux quiescent (Juarez *et al*., 2017; Fargier *et al*., 2012). High concentrations of electrophilic metabolites may override the suppression activity of MexS leading to conformational changes in MexT that activate efflux and return the cell to a redox balanced state (Fargier *et al*., 2012; Housseini B Issa, Phan and Broutin, 2018). Antibiotic resistance can arise through enhanced efflux as a result of mutations in *mexS* (Quick *et al*., 2014; Sobel, Neshat and Poole, 2005; Richardot *et al*., 2016) that prevent suppression of MexEF-OprN, or through activating mutations in *mexT* (Juarez *et al*., 2018), which result in constitutive expression of MexEF-OprN (Figure 1). These mutations represent genetic lesions that disrupt the native control cascade of this global regulatory system and come at a cost. The resistant cell now has reduced toxin degradation ability, reduced biofilm formation and reduced virulence associated with constitutive efflux (the ‘P1’ phenotype in PAO1), compared to the antibiotic susceptible phenotype with reduced efflux and enhanced virulence (the ‘P2’ phenotype in PAO1) (Fargier *et al*., 2012; Klockgether *et al*., 2010b; Tian *et al*., 2009b; Luong *et al*., 2014; Maseda *et al*., 2000b).

**Figure 1:**
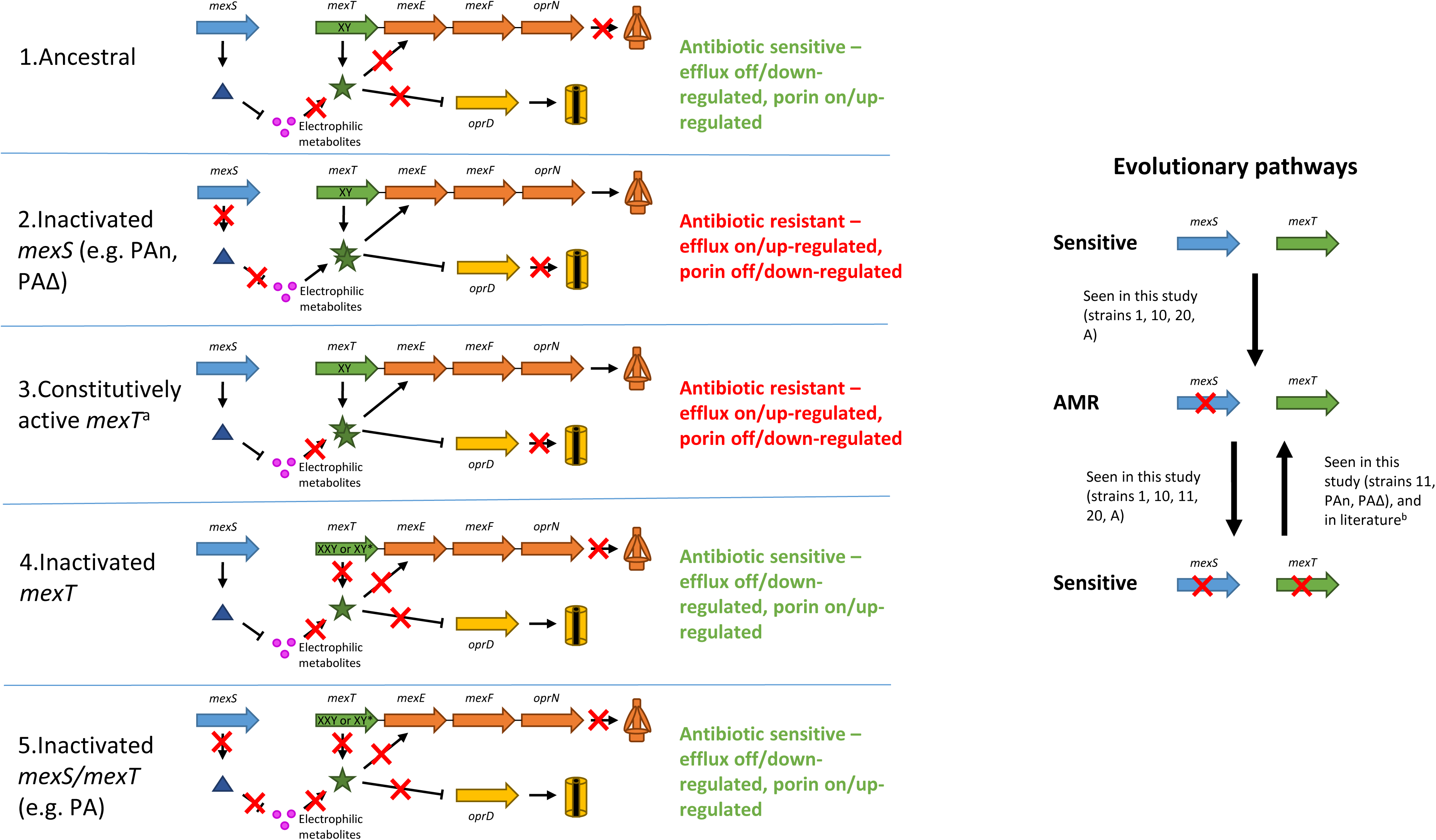
The impacts of mutations in mexS and mexT on efflux and porin production (left), and the direction of mutational changes observed in isolates in this study or in the literature (right). MexS/mexS are blue, MexT/mexT are green, the genes and proteins of the MexEF-OprN efflux system are orange, and the genes and proteins of the OprD porin are yellow. ^a^Juarez et al (2018) proposed that while MexT normally requires a cognate co-inducer to bind the inactive monomers to form the active oligomer, mutations in mexT produced MexT that spontaneously oligomerised into the active form. ^b^Maseda et al (2000a).

Investigation of the sub-lineages of the common research strain PAO1 revealed mutations in *mexS* and *mexT* that are rare in the wider *P. aeruginosa* population (Supplementary Table S2, Supplementary Figure S2)(Klockgether *et al*., 2010a; Stover *et al*., 2000; Luong *et al*., 2014; Sidorenko, Jatsenko and Kivisaar, 2017; LoVullo and Schweizer, 2020). Given that *mexT* is a global transcriptional regulator and is involved in both AMR and virulence, we investigated why disruptive genomic lesions arise so frequently in PAO1 *mexT* and described their phenotypic impact. To understand the impact of different genomic lesions, at a single locus, on the generation and cost of antibiotic resistance, we compared the transcriptomic and phenotypic effects of active versus inactive MexT in PAO1. We further explored our hypothesis that inactivation of MexT is a compensatory response to prior antibiotic resistance by selecting resistance in clinical isolates of *P. aeruginosa* and monitoring genetic changes in *mexT* after several generations of growth in antibiotic-free media. Growth conditions enabling reversion of antibiotic-resistant strains back to an antibiotic susceptible phenotype and genotype were identified, and explain the well-known mutations known to cause genome diversity in the reference strain, PAO1. Finally, we examine mutations in clinical isolates, in response to chloramphenicol treatment and explore the population level heterogenicity that is key to understanding the dynamics and diversity of adaption.

## Results and Discussion

### Inactivation of efflux via MexT in PAO1

*P. aeruginosa* PAO1 was isolated from a wound in 1954 by the Bruce Holloway laboratory, and in 2005 became the first sequenced *P. aeruginosa* strain and a designated reference (Stover *et al*., 2000; Holloway, 1955). However, the sequence of the global regulator *mexT* varies across PAO1 lineages, compared to the wider *P. aeruginosa* population, with an impact on phenotype (Table 1, Supplementary Table S2) (Klockgether *et al*., 2010a; LoVullo and Schweizer, 2020). Ancestral *mexT* has two imperfect repeats (‘X’: CGGCCA<u>G</u>C, and ‘Y’: CGGCCA<u>T</u>C, together defined as XY) but in the PAO1-UW reference sequence, duplication of the first 8-bp repeat (‘X’) transforms the ‘XY’ sequence to ‘XXY’ (Figure 2a, Supplementary Figure S3). This results in a reversible, inactivating frameshift (Maseda *et al*., 2000a) that was originally masked by mis-assignment of an upstream start codon in the PAO1-UW reference strain (LoVullo, E. D. and Schweizer, H. P. (2020)). Other types of inactivating mutations are also found in PAO1 *mexT* (referred to here as ‘XY*-type’ mutations) (Maseda et al., 2000a) and overall these changes block MexEF-OprN-mediated efflux and produce an antibiotic-sensitive strain (Supplementary Table S2). Mutations in the MexF component of the MexEF-OprN efflux pump (Luong *et al*., 2014) or other genes impacting efflux (Cai *et al*., 2023) can also arise are also observed in PAO1 and have a similar phenotypic impact, suggesting a common, adaptive benefit associated with reduced efflux. However, the increased frequency of MexT versus MexEF mutations suggests additional benefits from pleiotropic changes to global regulation mediated by MexT (Oshri *et al*., 2018). Conversely, *mexT* and *mexF* in non-PAO1 isolates within the sequence databases (NCBI, last accessed January 2021) were by a large majority intact (i.e. 98% were XY for *mexT* and 91% of *mexF* sequences lacked disruptive mutations), encoding an inducible efflux system, and highlighting the atypical nature of the PAO1 reference that is used routinely as a model.

**Figure 2.**
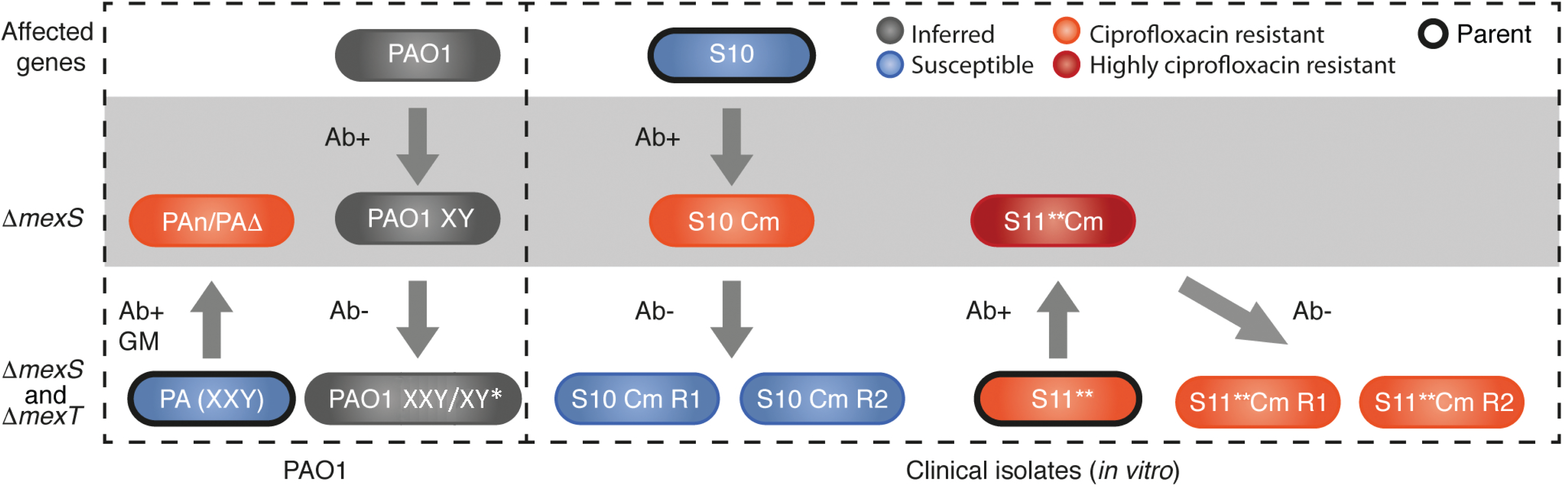
*mexT* sequence variation and evolutionary pathway for antibiotic resistance. a). The ancestral sequence of *mexT* is found in the XY formation. *P. aeruginosa* PAO1 *mexT* is disrupted by an 8 bp repeat (XXY). Other mutations (including those outside of this region) in *mexT* can also disrupt MexT (XY*). Numbering is according to the reference strain PAO1. b). Hypothesis for MexT disruption to switch off efflux under antibiotic-free conditions when MexS suppression has been lost. Ciprofloxacin resistance (≥ 2 µg/mL) typically emerges through disruption of *mexS* and reversion to ciprofloxacin sensitivity (< 2 µg/mL) emerges through disruption of *mexT*. Ab+; antibiotic present, Ab-; antibiotic-free, the P1 and P2 phenotypes are defined by Luong *et al*., 2014 and the P2 phenotype displays increased virulence.

**Table 1.** Expected characteristics of *P. aeruginosa* isolates and mutations present. *P. aeruginosa* strains UCBPP-PA14, mPAO1/P1, PAO1 (Reference), PA1R and C7447m are named according to the *Pseudomonas* Genome Database. UCBPP-PA14 is proposed to represent native control over efflux in *P. aeruginosa*. <u>Key</u> +: gene is expected to be expressed producing functional gene product, -: gene expression is not expected to lead to a functional gene product; Cip^S^: ciprofloxacin susceptible; Cip^R^: ciprofloxacin resistant, P1/P2: phenotypes proposed by Luong *et al*., 2014 (P1; slow growing, lower virulence phenotype, antibiotic resistant P2; fast growing, higher virulence phenotype, antibiotic sensitive).

| Genotype |  | Expected phenotype |  |  | Representative *P.aeruginosa* strain (*mexT* genotype) |
| --- | --- | --- | --- | --- | --- |
| *mexS* | *mexT* | Classification | Efflux | Resistance |  |
| + | + | P2 | off | Cip<sup>S</sup> | UCBPP-PA14 (*mexT* XY) |
| - | + | P1 | on | Cip<sup>R</sup> | mPAO1/P1 (*mexT* XY) |
| - | - | P2 | off | Cip<sup>S</sup> | PAO1 (Reference) (*mexT* XXY) |
| + | - | P2 | off | Cip<sup>S</sup> | PA1R/C7447m (*mexT* XXY) |

### Disruption of MexT is not restricted to PAO1

The *mexT* XXY form (Supplementary Figure 3) is only described for PAO1 isolates but this genotype is not limited to the reference strain (Supplementary Table S2). To assess the prevalence of these variants in the general *P. aeruginosa* population, we reviewed all sequences deposited in the NCBI database (accessed March 2018) and identified rare 8-bp insertions (XXY) in *mexT* including within a mucoid cystic fibrosis *P. aeruginosa* C7447m isolate (Yin *et al*., 2013) and a phage-resistant laboratory derivative of the respiratory isolate *P. aeruginosa* PA1 (Lu *et al*., 2015; Le *et al*., 2014). Modification or deletion of *mexT* has also been reported for lung-adapted clinical isolates (Warren *et al*., 2011; Smith *et al*., 2006; Hilliam *et al*., 2017; La Rosa, Johansen and Molin, 2018). This confirmed that overall *mexT* was well conserved and that changes from the wild-type XY form of *mexT* to the *mexT* XXY genotype exist but are rare, meaning the vast majority of studied strains differ from the PAO1-UW reference strain.

### Refunctionalisation of MexT in PAO1 produces an antibiotic-resistant phenotype

Resistance to ciprofloxacin in *P. aeruginosa* can emerge through mutations in MexS, which activates MexT and upregulates MexEF-OprN-mediated efflux (Figure 1). Although efflux enables survival in the presence of antibiotics, it may be maladaptive in the absence of antibiotics. We hypothesised that MexT disruption is a strategy to switch off efflux (and the associated stress response) under antibiotic-free conditions when MexS suppression has been lost during the acquisition of antibiotic resistance (Figure 2b), and that this is what has occurred to produce the antibiotic-sensitive PAO1-UW reference strain (encoding inactive MexS-N249D and *mexT*-XXY). To address this, we established the phenotypic impact of reintroducing functional (XY) MexT into a PAO1 MexS-null background.

We used a strain of PAO1 with a MexS N249D null mutation (Richardot *et al*., 2016) and the inactive XXY formation of MexT (named strain PA) to construct an isogenic mutant (PAΔ) with one X repeat deleted (to produce an XY genotype) (Figure 2a, Figure 3). We also subjected PA to ciprofloxacin selection pressure and isolated an *nfxC*-type mutant (PAn) containing a single *mexT* X repeat (Köhler *et al*., 2001)(Supplementary Table S1). Deletion of the second 8 bp repeat in *mexT* restored the correct reading frame in PAΔ and PAn, resulting in *mexT* expression increasing by over 1.6 log_2_ fold compared to PA (p < 0.001, Supplementary Dataset S1). Functionalisation of MexT in PAΔ and PAn produced strains resistant to chloramphenicol, ciprofloxacin and imipenem consistent with overexpression of the MexEF-OprN pump in the absence of functional MexS (Figure 1, Supplementary Dataset S1 and S2). This indicates that antibiotic-resistant PAO1 with inactive MexS but functional MexT (XY genotype) is the likely ancestor to the sensitive PAO1 strains with inactive MexS and MexT, where MexT disruptions have arisen through various mechanisms (XY* and XXY) (Figure 1, Figure 3). After our study was completed, separate research (Lee, Gallagher and Manoil, 2021) was published suggesting the same evolutionary trajectory. Here, the authors restored the ancestral (functional) MexS protein in the PAO1 reference strain and highlighted the salt-sensitivity associated with the MexS-null mutant widely used in current research as well as the atypical nature of PAO1.

**Figure 3.**
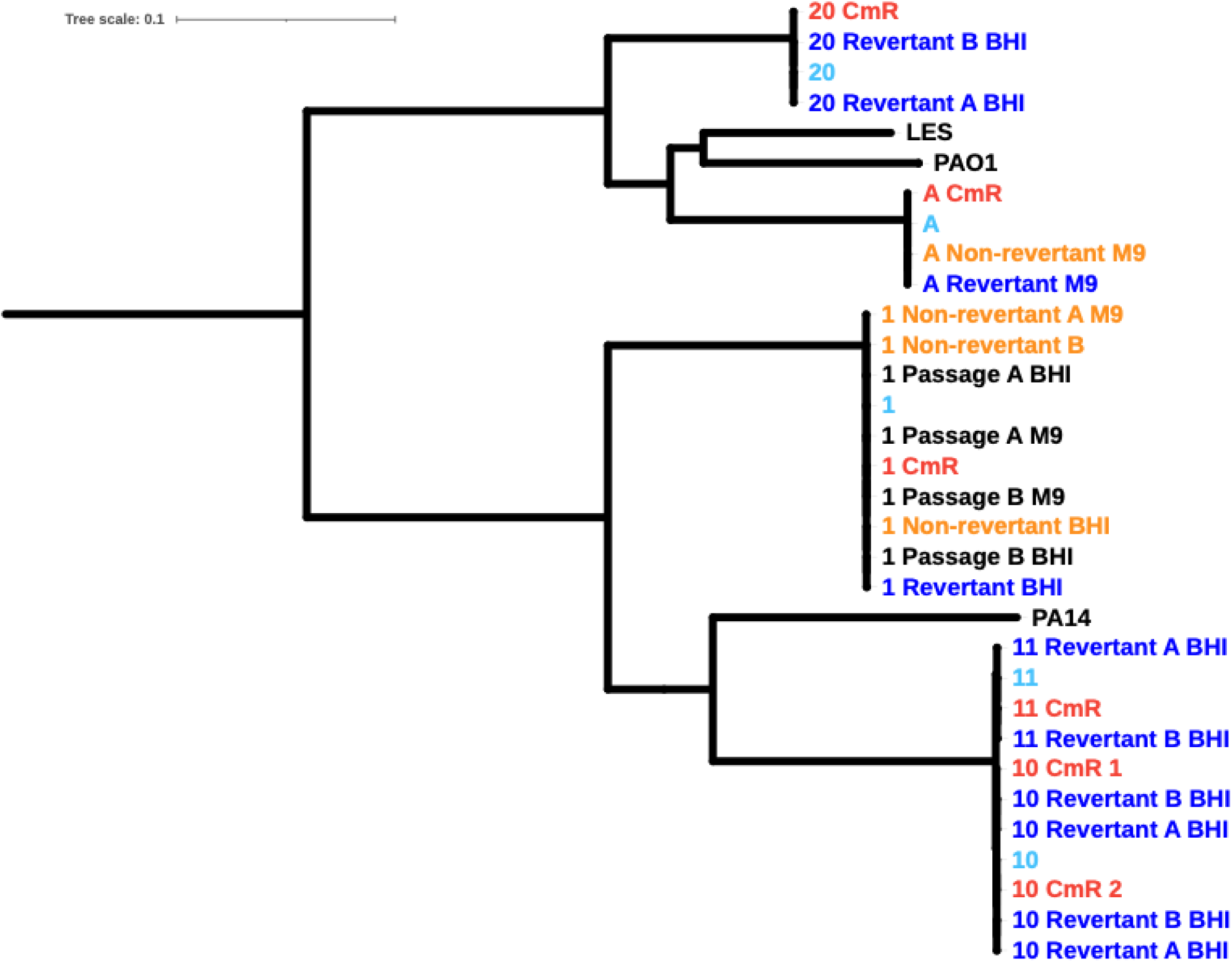
*in vitro* evolution of antibiotic resistance in P. aeruginosa. Impact of addition and removal of antibiotic selection pressure upon phenotypes and genotypes in *P. aeruginosa* isolates. Dashed line box on the left indicates responses of PAO1 isolates (including PAΔ, generated through genetic modification: GM). Dashed line box on the right indicates responses of clinical isolates. Colours denote susceptibility (blue, < 2 µg/mL) and resistance (red, ≥ 2 µg/mL) to ciprofloxacin; high level resistance to ciprofloxacin (≥ 32 µg/mL) was conferred by mutations in *gyrA* (**), denoted by dark red. Phenotypes of PAO1 isolates in grey are inferred. Ab+: chloramphenicol selection; Ab-: antibiotic-free; Cm: chloramphenicol-resistant mutant; R: chloramphenicol-sensitive revertant.

### Antibiotic exposure selects for resistance via inactivating MexS

We have shown that in the PAO1 inactive MexS/MexT background, the route to antibiotic resistance is through refunctionalising MexT. However, most *P. aeruginosa* isolates have the ancestral active MexS/MexT background (Figure 1), and previous work has shown that inactivating MexS results in an antibiotic resistant phenotype (Richardot *et al*., 2016; Lee, Gallagher and Manoil, 2021; Kostylev et al., 2023). To characterise the involvement of MexS and MexT in the evolution of antibiotic resistance across varied genotypic backgrounds, we exposed *P. aeruginosa* clinical strains to chloramphenicol and selected isolates with increased resistance (400 µg/mL). A total of 40 mutants were picked from 33 *P. aeruginosa* genotypes (one or two mutants per parent strain). At the end of the evolution experiments, whole genome sequencing was performed on six selected mutants, generated from five distinct parent strains, to confirm the mechanism of chloramphenicol resistance. This also enabled us to place these strains within a wider phylogeny of *P. aeruginosa* (Figure 4). Sequencing revealed that *in vitro* evolution of chloramphenicol resistance was associated with *mexS* truncations or frame shifts in 5/6 cases, consistent with the literature (Table 2). In the remaining case of Strain 11, the parent likely had an inactive MexS/MexT background, similar to PAO1 whereby resistance instead emerged through refunctionalisation of MexT. Consistent with this, chloramphenicol selection in Strain 11 produced a mutant with a 1-bp deletion in *mexT*, creating a frameshift and producing an active protein. Notably, the MexS mutation seen in the ciprofloxacin-resistant clinical isolate Strain 11 (Table 2, Figure 1) was also observed *in vitro* (A27fs in strain A) confirming the relevance of the *in vitro* resistance mechanisms to *in vivo* resistance mechanisms. New mutations in topoisomerases/gyrases were not observed, consistent with chloramphenicol rather than ciprofloxacin selection (Bruchmann *et al*., 2013). Changes were not detected in *cmrA* by ARIBA analysis, which has also been associated with MexEF-OprN-mediated resistance (Juarez *et al*., 2017). In all six sequenced mutants, MIC values for ciprofloxacin increased alongside an increase in chloramphenicol MIC, and in 4/6 mutants the imipenem MIC also increased (Table 2), in line with the known function of MexEF-OprN and efflux of chloramphenicol and ciprofloxacin and import of imipenem (Köhler *et al*., 2001). Of note, the two mutants where imipenem MIC did not increase (Strain 10 Cm^R^ 1 and Strain 10 Cm^R^ 2) contained a frameshift mutation in *oprD* in the parent strain and so were already resistant to imipenem. Taken together, these data show that in the active MexS/MexT background, the predominant mechanism through which *P. aeruginosa* switches from an antibiotic sensitive to resistant phenotype is through mutation of *mexS*, resulting in active MexT.

**Figure 4.**
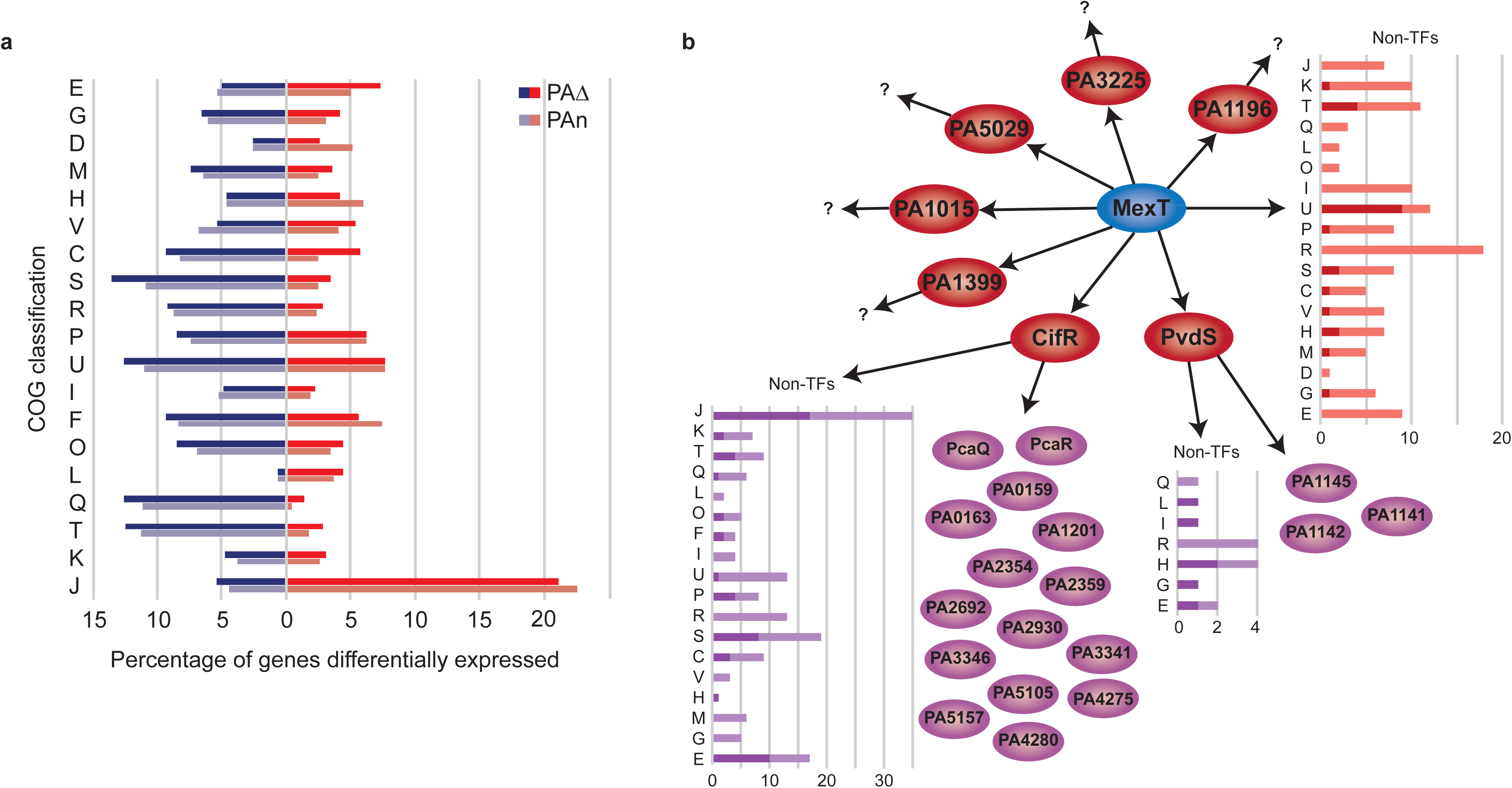
Phylogenetic analysis of *P. aeruginosa* isolates. Core SNP tree of *P. aeruginosa* parent strains (light blue), *in vitro*-generated chloramphenicol resistant mutants (CmR, red), passaged chloramphenicol sensitive revertant (dark blue) or resistant non-revertant strains (orange). Reference strains including *P. aeruginosa* PAO1-UW, *P. aeruginosa* UCBPP-PA14 and *P. aeruginosa* LESB58 are also provided for comparison.

**Table 2.**
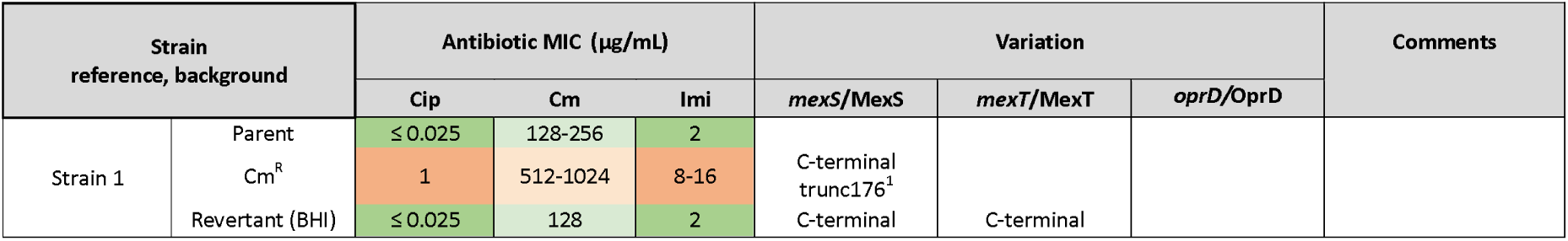
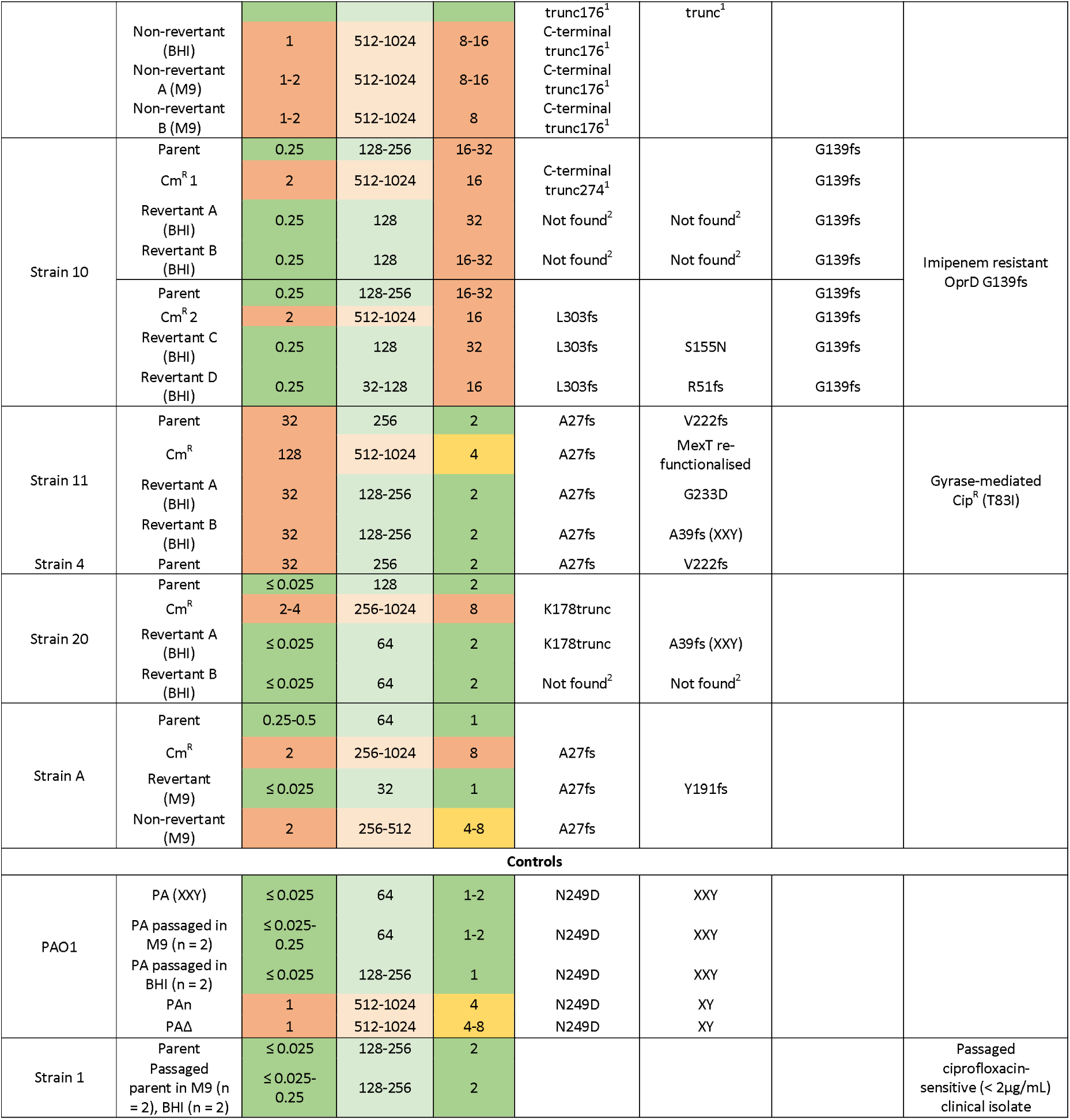
Antibiotic susceptibility and genotype of *P. aeruginosa* strains evolved in this study (MIC range, n =2). The genotype was determined by analysis of the raw sequence reads using the ARIBA tool as described in the methods, protein sequences were also extracted by BLAST for confirmation of changes as described in the methods. The clinical breakpoint at which strains are considered to be resistant is 1 µg/mL for ciprofloxacin (cip), and 8 µg/mL for imipenem (imi)^1^. Cm: chloramphenicol, CmR: chloramphenicol resistance; trunc: truncation; fs: frameshift, BHI: passage in Brain Heart Infusion broth, M9: passage in SSM9PR broth, XY: ancestral *mexT* with two imperfect ‘XY’ repeats, XXY: disrupted *mexT* with duplication of the first 8bp repeat, ^1^ results obtained using protein sequences extracted by BLAST, ^2^ sequences not found by ARIBA analysis or extraction by BLAST.

### Global transcriptomic impact of MexT activation

In addition to its role in antibiotic resistance, MexT is known to be involved in the regulation of a number of other genes (Tian *et al*., 2009a). In order to determine the wider impact on transcription in the antibiotic-sensitive MexT-inactive phenotype (XXY) and the antibiotic-resistant MexT-active phenotype (XY), we performed transcriptomics on strains PAΔ and PAn (carrying active MexT) and their parental strain PA (with inactive MexT) (Figure 1, Figure 3). First, we sequenced the PA parent strain and both PAΔ and PAn variants on the Illumina platform to confirm the repeat region in *mexT* to be XY in both PAΔ and PAn. Thus, the transcriptional effects we describe below are due to changes in MexT, relative to the parent PA.

Globally, with respect to PA, 600 genes in PAΔ showed a reduction in expression (log_2_FC < -1 and p < 0.05) and 327 showed increased expression (log_2_FC > 1 and p < 0.05), meaning that the change in *mexT* affected expression of 1/6 of the genome (922/5573 genes) with only a slightly reduced impact for PAn (806/5573). Despite the close genetic similarity of the variants, only 80% of the same genes were down-regulated in both (507/612), with even fewer of the up-regulated genes in common (68%; 245/358). Genetic differences and stochastic effects may account for this variation but comparison to PA allowed us to identify common genes under the control of MexT.

Using COG annotations from the *Pseudomonas* Genome Database (Winsor *et al*., 2016), we assessed which functional categories were most affected in PAΔ and PAn (Figure 5a). The effects were wide ranging but proportionally, the largest increase in expression was seen in translation, ribosomal structure and biogenesis (category J), confirming the important regulatory role of *mexT*. Over 10% of genes in four categories were down-regulated: secretion (U), signal transduction (T), secondary metabolites (Q) and function unknown (S) suggesting that although *mexT* is an activator for MexEF-OprN, it also enables repression of certain characteristics possibly by indirect control via other transcriptional regulators (Supplementary Dataset S1 and S2), which has also been reported for other LysR regulators (Vercammen *et al*., 2015).

**Figure 5.**
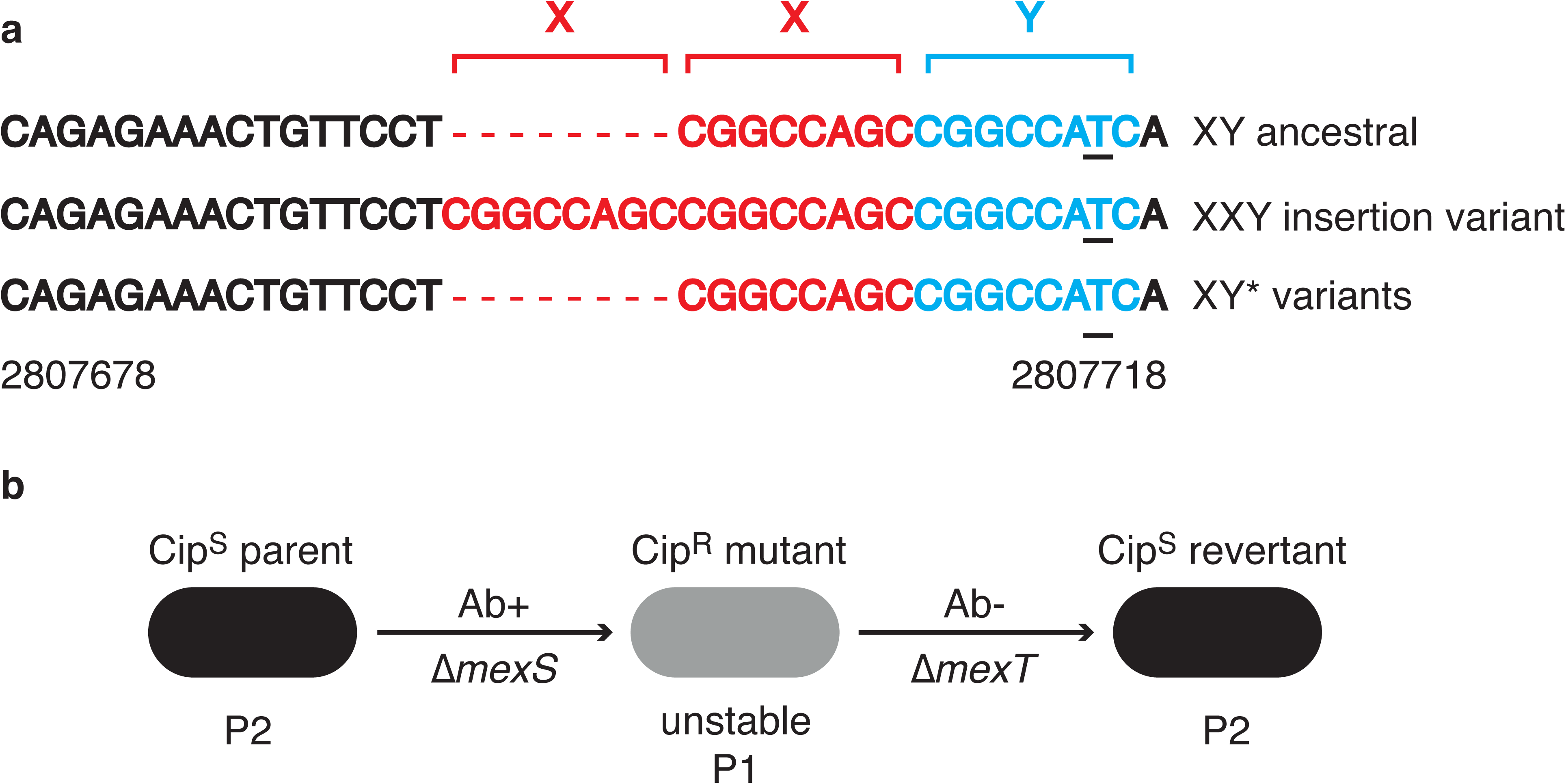
Impact of MexT on gene expression and regulation in PAO1. a). Differentially expressed genes in PAn and PAΔ (active MexT XY) compared to the parent PA (inactive MexT XXY). Bars indicate the percentage of the genome total for each COG category. Blue = down-regulated; red = up-regulated. Absolute numbers per category are in Supporting Information. b). Transcriptional regulatory network of MexT. Transcription factors (TFs) in red are directly affected by MexT. Downstream effects are shown in purple where known; TFs in purple and non-TFs in bar charts. Numbers of non-TFs are displayed by COG category, with bold colour indicating how many genes were found to be differentially expressed in PAΔ or PAn, compared to PA. E: amino acid transport and metabolism; G: carbohydrate transport and metabolism; D: cell cycle control, cell division, chromosome partitioning; M: cell wall/membrane/envelope biogenesis; H: coenzyme transport and metabolism; V: defence mechanisms; C: energy production and conversion; S: function unknown; R: general function prediction only; P: inorganic ion transport and metabolism; U: intracellular trafficking, secretion, and vesicular transport; I: lipid transport and metabolism; F: nucleotide transport and metabolism; O: post-translational modification, protein turnover, chaperones; L: replication, recombination and repair; Q: secondary metabolites biosynthesis, transport and catabolism; T: signal transduction mechanisms; K: transcription; J: translation, ribosomal structure and biogenesis.

Transcriptionally active MexT (in PAΔ and PAn) had the greatest impact upon expression of *mexEF* and *oprN* in both variants (log_2_FC > 7, p < 0.001 for all 3 genes), as expected given its known role regulating the MexEF-OprN efflux pump, as well as hypothetical signal peptides (Supplementary Dataset S1 and S2) reportedly under the control of MexT (Vercammen *et al*., 2015; Tian *et al*., 2009a). In line with the literature, down-regulation of virulence genes involved in phenazine, rhamnolipid and hydrogen cyanide production (Köhler *et al*., 2001; Tian *et al*., 2009b;) was observed as well as partial upregulation of the nitrogen respiratory chain (nitrite genes) for PAΔ, which is a reported compensation strategy for efflux in *P. aeruginosa* (Olivares, Álvarez-Ortega and Martinez, 2014). In contrast to a previous report of a *mexT*-controlled *mexEF* over-expresser (Linares *et al*., 2005), the T3SS encoded by *pscB*-*L* was up-regulated, along with its activator *exsA*. Functional MexT also activated detoxification gene sets (Fetar *et al*., 2011) including the nitrosative response (PA3229 and PA4881, log_2_FC >5.5, p < 0.001), *xenB* operon (PA4354-56, log_2_FC ≥ 1.4, p < 0.001) and several ABC transporters (PA2811-2, log_2_FC ≥ 1.9, p < 0.001), consistent with a possible role for MexEF-OprN in cellular protection during host colonisation (Fetar *et al*., 2011). The oxidoreductase *mexS* (PA2491), implicated in regulation of the MexEF-OprN efflux pump, was also upregulated, consistent with active MexT and MexEF, and promoting the primary mechanism of detoxification (log_2_FC ∼ 3, p < 0.001) (Fargier *et al*., 2012). However, since a null mutation is present in PA *mexS* (N249D), transcription of *mexS* does not equate to a functional MexS protein (Richardot *et al*., 2016; Lee, Gallagher and Manoil, 2021).

Toxin export through MexEF-OprN is a proposed ‘back-up’ for cell survival, but may also serve to divert energy from virulence traits during cellular stress by export of quorum sensing molecules (Lamarche and Déziel, 2011; Kim, Yoon and Choi, 2015; Kumar and Schweizer, 2011). Altered efflux has been reported for PAO1 in rat tissue and *in vivo* models and may be either active (Join-Lambert *et al*., 2001) or inactive (Frisk *et al*., 2004; Luong *et al*., 2014) suggesting that niche and temporal responsiveness is key. PAO1 strains that have lost regulation of efflux may rely on alternative strategies for control of export, potentially at the expense of systems related to virulence.

### Regulatory network demonstrates a route to altered virulence

*P. aeruginosa* possesses two quorum-sensing systems, *las* and *rhl,* which are important for acute virulence (Shrout *et al*., 2006). Both *lasR* and *rhlR* were down-regulated in the antibiotic-resistant mutants and may explain reduced swimming and swarming seen in these strains, indicative of reduced virulence (Winstanley, O’Brien and Brockhurst, 2016) (Supplementary Figure S1). While a considerable number of other genes were found to be either up- or down-regulated in response to active versus inactive MexT, the relationship of MexT to these genes was not known. In order to investigate which genes are regulated by MexT, the transcriptional regulatory network downstream of MexT (Figure 5b) was reconstructed using published MexT binding sites (Maseda, Uwate and Nakae, 2010; Tian *et al*., 2009a). At the top level (direct regulation), MexT binding sites were associated with 166 genes, of which 7 were transcription factors (TFs)(Supplementary Dataset S3). Of these seven, two TFs (CifR and PvdS) had known TF binding sites (Ballok *et al*., 2012; Wilson, McMorran and Lamont, 2001; Ochsner *et al*., 2002), which we used to determine secondary, indirect regulation. This revealed that MexT directly and indirectly (particularly through CifR) mediates a wide range of functions (based on COG classification, Figure 5b). The identification of five MexT-regulated TFs with unknown binding sites provides scope for future study into the regulatory cascades that cause the observed gene expression changes and phenotypes.

Overlaying our RNA-seq data onto this regulatory network indicated that functionalisation of *mexT* significantly affected one of the seven TFs, *pvdS* (log_2_FC of ∼0.8 in PAΔ/PAn, p < 0.001), a sigma factor within the *pvd* locus. In PAΔ and PAn, upregulation of siderophore (pyoverdine) expression via PvdS (e.g. *pvdEFHNPT* Log2FC *≥ 1*, p < 0.001) (Miyazaki *et al*., 1995) correlated with reduced phenazine expression. Phenazines are redox agents with a role in iron metabolism. Both metabolites use ferric iron as a substrate but have complementary roles: phenazines generate bioavailable ferrous iron from sources of ferric iron (Wang *et al*., 2011) and pyoverdines sequester ferric iron directly (Cornelis and Dingemans, 2013). Active MexT in PAn and PAΔ reduced ferrous iron metabolism (phenazine biosynthesis and the ferrous iron transporter *feoAB*) and increased pyoverdine production (involved in ferric iron uptake) as well as genes linked to the acquisition of haem iron (*phuRT*). Reduced phenazine gene cluster expression in PAΔ and PAn suggests a reduced virulence phenotype (Panayidou *et al*., 2020). It is conceivable that active MexT and active efflux, as in PAΔ and PAn, is associated with colonisation of clinically relevant sites and expression of appropriate iron utilisation genes (Cornelis and Dingemans, 2013), whereas inactive MexT is associated with virulence and infection. For example, bioavailable ferrous iron is present in the infected lung (Cornelis and Dingemans, 2013) and the pyoverdine route to iron acquisition is often lost during chronic infection (Hilliam *et al*., 2017), whereas production of redox-active phenazines is key to virulence (Lau *et al*., 2004). Similarly, in a rat intestinal model the frequency of *P. aeruginosa* carrying inactive MexT was high at the site of surgical trauma and low in isolates colonising a distant healthy intestinal site (Olivas *et al*., 2012). Overall, these data show that regulation of MexS and MexT by *P. aeruginosa* can be used to switch between a more benign but drug-resistant phenotype and a virulent but more drug-sensitive phenotype. This regulation enables the bacteria to be responsive to niche-specific stressors, signalling molecules and to balance appropriate substrate import against virulence. However, our understanding of the phenotypic range and regulatory control over (potentially independent) efflux and other MexT-mediated global effects remains poorly understood (Schweizer, 2003). Following our studies, we noted that Kostylev et al., (2023) came to similar conclusions around the impact of mutated MexS in PAO1 on the activity of MexT and efflux and the consequences for virulence, in particular regarding quorum sensing via RhlI-R.

Some phenazines, such as pyocyanin, are toxins that are important for the establishment of *P. aeruginosa* infection and virulence (Lau *et al*., 2004). Therefore, we tested the virulence of PA, PAΔ and PAn in a *Galleria melonella* infection model. Strain PAΔ, with active MexT, was significantly less virulent than the parental PA strain, with inactive MexT (p*=3.27e-7*). PAn (with active MexT) was also less virulent, but not significantly different compared to PA. However, the efflux-active mutants (PAΔ and PAn) also differed significantly from each other in terms of virulence (p*=4.02e-4)* indicating the role of other factors. Data from a previous study has shown that production of virulence factors was inversely correlated with *mexE* expression (Richardot *et al*., 2016). A reduced virulence phenotype of MexT-active strains may be partly explained by the down-regulation of phenazine biosynthesis (Cezairliyan *et al*., 2013), (Supplementary Figure S1, D).

### Changes in MexT cause a metabolic shift in substrate utilisation

In addition to the role of MexT in directly and indirectly regulating genes linked to antibiotic resistance and virulence, we hypothesised that the large changes in global gene expression seen in strains PAΔ and PAn (active MexT) compared with their parental strain PA (inactive MexT from XXY genotype) would have a wider impact on bacterial metabolism. To investigate this, we carried out metabolic phenotyping of these three strains. As a logical consequence of reduced global transcription, metabolic phenotyping of 626 compounds using the Omnilog system indicated that PAΔ and PAn were less metabolically active than PA. Both PAΔ and PAn were particularly defective in protein, amino acid and nucleoside compound metabolism (Supplementary Dataset S4). These processes are energy (ATP)-expensive, and their reduction may help balance the increased efflux, which is also ATP-expensive. Overexpression of *mexEF-oprN* is linked to decreased amounts of the outer membrane porin OprD, a mechanism by which basic amino acids and peptides are transported into the cell (Ochs *et al*., 1999). We observed a 0.8-1.2 log_2_ fold reduction in *oprD* expression in PAΔ and PAn (p ≤ 0.012, Supplementary Dataset S1) suggesting a genotype-phenotype link, as well as differences in utilisation of short peptides. Other transport systems such as the branched chain amino acid transport (*bra*) operon were upregulated in PAΔ and PAn and may be compensating for the reduction in *oprD* expression, as well as the reduction in protein, amino acid and nucleoside compound metabolism by taking up resources from the growth medium. Differences were not observed in nitrite utilisation in PAΔ and PAn despite upregulation of the nitrite genes.

The shift away from central metabolism in antibiotic-resistant strains is logical since efflux is part of the stress response. In the absence of host inducers or antibiotics, quiescent MexEF-OprN may offer a fitness benefit. This leads us to speculate that nutrient availability, as found in rich laboratory media or relatively amino acid rich host environments such as the lung (Winstanley, O’Brien and Brockhurst, 2016), could select for the higher metabolic activity of strains carrying disrupted *mexT* in order to exploit the available energy sources. For example, phosphate availability has already been linked to the activity of MexT whereby phosphate supplementation prevented MexT disruption in a rat intestinal model (Olivas *et al*., 2012). Growth of *P. aeruginosa* in peptidic laboratory media may require secreted proteases, a product of quorum sensing (Oshri *et al*., 2018), which is deficient in these efflux-positive strains (Supplementary Dataset S1). Indeed, several recent studies in PAO1 suggest that MexT is disrupted to promote RhlR-mediated quorum sensing in LasR-null strains enabling growth in peptide-based media (Oshri *et al*., 2018; Kostylev *et al*., 2019; Cheng *et al*., 2022) or to bypass the stringent response and produce virulent strains (Figueroa et al., 2025), although we propose *mexS* rather than *lasR* or other factors as a driving force. However, the impact in a minimal medium remains to be tested and further work is needed to determine the impact of LasR versus MexS mutations in driving shutdown of efflux in the atypical PAO1 background. As shown above, this switch to a more metabolically active phenotype via MexT inactivation, is also associated with reduced antibiotic resistance but increased virulence.

Overall, metabolic profiling revealed a profound phenotypic switch associated with active or inactive MexT in PAO1. However, the impact of mutations in *mexS* is unknown. In strains PAΔ and PAn, efflux was switched on by reactivation of MexT and similar observations have been made by others (Juarez *et al*., 2018). However, in some strains the phenotypic switch is governed by the inactivation of MexS rather than (re)activation of MexT (Figure 1). Future work characterising the transcriptomes of our mutant collection (Table 2) could unravel the impact of individual *mexS*, *mexT* and *mexEF-OprN* mutations on *P. aeruginosa* physiology.

### Reverting from antibiotic resistance to antibiotic sensitivity is achieved through disruption of MexT

We have observed the switch from an antibiotic-sensitive to an antibiotic-resistant phenotype in both our PAO1-derived strains (PAΔ and PAn) and by passaging our selected clinical isolates on antibiotic media (Table 2). However, this switch to an antibiotic-resistant phenotype (non-functional MexS) has arisen from two different genetic backgrounds – either an antibiotic-sensitive non-functional MexS/MexT background (strains PA and 11) or an antibiotic sensitive functional MexS/MexT background (strains 10, 20, A) (Figure 1). This raises the question: in the absence of antibiotic selective pressure, do the bacteria revert to sensitivity and, if so, do they do this by refunctionalising MexS or by inactivating MexT? In order to answer this, we passaged our strains, previously selected for increased chloramphenicol resistance, in the absence of antibiotic, and sequenced the genomes of isolates that reverted to sensitivity. Analysis of the chloramphenicol-sensitive revertants confirmed our hypothesis that antibiotic resistance can be an unstable phenotype during laboratory passage and that MexT is disrupted to switch off efflux and restore sensitivity (Figure 3, Figure 1). We found that MexT was inactivated e.g. by indels or nonsynonymous mutations and, in some cases, multiple strategies for inactivation were observed within a single population (Table 2). Significantly, the frameshift in clinical strain 20 was due to an 8-bp insertion producing the *mexT* XXY form seen in PAO1, consistent with our evolutionary proposal that variation in PAO1 *mexT* XXY is a consequence of antibiotic resistance and subsequent reversion to sensitivity following laboratory culture. Indeed, this study demonstrates that revertants can be readily obtained following standard laboratory growth over relatively few passages. In some revertant strains, *mexT* was not detected. Complete deletion of MexT has previously been identified in strains isolated from the lung (Warren *et al*., 2011; Smith *et al*., 2006; Hilliam *et al*., 2017). Mutations arose rapidly in *mexT* (within 6 passages) and these mutants appeared to be dominant as they were detected by screening fewer than ten colonies (from cultures with reduced growth on antibiotic), implying a fitness benefit since selection was based solely on enhanced growth during passage. Disruption of MexT to revert to an antibiotic-sensitive phenotype is not limited to strains closely related to the reference PAO1 and was observed in isolates representing all major branches of our *P. aeruginosa* phylogenetic tree (Figure 4). SNP analysis was performed to assess genome-wide changes that could also account for altered antibiotic resistance (Supplementary Dataset S5). However, no other changes were consistently identified across the evolved strain sets, confirming that disruption of MexT is key to reversion. We initially hypothesised that antibiotic-resistant strains reverted to an antibiotic-sensitive phenotype in the absence of antibiotic selective pressure since downregulation of MexEF-OprN-mediated efflux increases expression of the inversely co-regulated OprD porin (Figure 1) involved in amino acid import to the cell. However, an imipenem-resistant strain lacking functional OprD was available (Strain 10, Table 2) and this strain also reverted to a susceptible phenotype through inactivation of MexT (Figure 3, Table 2), suggesting a limited role for OprD in the growth conditions used.

To determine whether mutations in *mexS* and *mexT* affect bacterial fitness we analysed bacterial growth of the ancestors, mutants and revertants for the antibiotic sensitive strain A and the imipenem-resistant Strain 10 (Table 2, Suppl Figure S4). Regardless of strain background, resistant mutants arising through inactivation of *mexS* performed better than the ancestor in the presence of antibiotic selective pressure, and worse than the ancestor in the absence of selective pressure (Suppl Figure S4). However, different *mexs*/*mexT* backgrounds of the revertants were associated with different growth characteristics. Complete deletion of *mexS* and *mexT* (Strain 10; revertants A and B) produced revertants that were better able to grow under selective pressure than the ancestor (though not quite as well as the resistant mutant) but in the absence of selective pressure out-perform the ancestor (Suppl Figure S4). Conversely, a frame-shifted MexS coupled with a single amino acid change in MexT (Strain 10; revertant C) resulted in no difference between the revertant and ancestor under selective pressure but significantly worse growth of the revertant in the absence of selection (Suppl Figure S4). Similar results were observed when both *mexS* and *mexT* are disrupted by a frameshift, though this revertant (Strain 10; revertant D) is less impaired compared to the ancestor in the absence of selection. Finally, the revertant of strain A which carried frameshift mutations in both MexS and MexT had comparable growth to the ancestor with and without selective pressure and these strains achieved the highest cell densities overall. While these results are specific to the growth conditions used in this study they indicate that the means by which strains mutate and revert to a sensitive phenotype impacts bacterial fitness, implying that different mechanisms of reversion may be selected under different environmental conditions or in different genetic backgrounds.

To investigate whether *mexS*/*mexT*-disrupted genotypes observed *in vitro* are found in clinical strains, we reviewed our clinical collection and identified two genetically-similar catheter urine isolates of *P. aeruginosa* with both *mexS* and *mexT* impaired by frame-shift mutations (Strain 4 and Strain 11, Figure 4, Table 2). It is possible that these mutations reflect antibiotic resistance through *mexS* and subsequent adaptation to virulence in a host environment by disruption of *mexT*, although mutations arising during diagnostic processing cannot be ruled out since we observed changes during laboratory passage. These isolates had high level ciprofloxacin-resistance (32 µg/mL) due to a T83I mutation in *gyrA* (Bruchmann *et al*., 2013). The order of mutation events is unknown, though previous work detected mutations in *gyrA* and *gyrB* far more frequently in genetic backgrounds already carrying mutations causing expression of efflux systems than in sensitive wild-type backgrounds (Llanes *et al*., 2011). If MexT inactivation occurred first, resistance through gyrase mutations upon subsequent antibiotic exposure would produce both a virulent and highly resistant strain.

In order to assess the impact of repeated antibiotic selection to reflect antibiotic cycles in the clinic, we used Strain 11, which already carried the T83I mutation in *gyrA* and had non-functional MexS and MexT. Using this ciprofloxacin-resistant clinical isolate, we selected for increased chloramphenicol resistance, which arose through a 1-bp deletion causing a frameshift that re-functionalised MexT and further increased ciprofloxacin resistance 4-fold (from 32 to 128 µg/mL, Table 2). Mutations in gyrases and increased efflux are known to be independent, additive mechanisms of fluoroquinolone resistance (Bruchmann *et al*., 2013) and were associated with the highest observed resistance. Upon antibiotic-free passage, disruption of MexT was observed as a nonsynonymous amino acid change (G233D, Revertant A) or an 8-bp insertion (MexT XXY, Revertant B), but the strain remained antibiotic resistant (reverting to its previous ciprofloxacin MIC of 32 µg/mL) due to the gyrase T83I background. The reversible introduction and excision of a short DNA sequence in *mexT* reflects the scenario seen with PAO1 XY versus XXY forms. This supports the proposed evolutionary pathway for PAO1 and demonstrates that once strains have acquired mutations in *mexS*, the switch between the antibiotic-resistant, less virulent phenotype and the antibiotic-sensitive, more virulent phenotype is achieved through inactivation of MexT rather than refunctionalisation of MexS (Figure 1). There are rare exceptions in PAO1 where MexT is disrupted whilst MexS appears intact (Supplementary Figure S2, PAO1 lineages PAO1_VE2, VE13 and H2O); we speculate that for PAO1 in these cases MexS has subsequently reverted to a functional form. Finally, SNP and SNPEff analysis was carried out for each set of parent and revertant strains across the whole genome in order to identify if mutations elsewhere in the genome could have been responsible for any effects seen. These results are presented in Supplementary Dataset S5 and Supplementary Figure S5 and, although no other key changes were noted, further work is required to understand the broader circumstances leading to disruption of MexT.

### Conclusions

*P. aeruginosa* pathogenesis is frequently modelled using the reference strain PAO1. However, laboratory lineages of PAO1 are phenotypically diverse due to variation in *mexT*, and also differ in *mexS* compared to other *P. aeruginosa* lineages. We propose that antibiotic resistance in *P. aeruginosa* PAO1, conferred by an ancestral *mexS* mutation, drives plasticity in *mexT* and this evolutionary pathway was demonstrated with clinical isolates *in vitro*, although the relevance to *in vivo* mechanisms is unknown. Overall, in ancestral antibiotic-sensitive *P. aeruginos*a (Figure 1), antibiotic resistance emerges through the loss of homeostatic control of efflux by inactivation of MexS. In these antibiotic-resistant *P. aeruginosa*, in addition to the loss of control over efflux, virulence and metabolism are also reduced unless mutations emerge inactivating the global regulator MexT. This restores antibiotic sensitivity and increases metabolism and production of virulence factors, but does not restore homeostatic control of efflux as inactive MexS cannot modulate MexT (Figure 1, inactivated *mexS/mexT*). Further antibiotic exposure may then cause the bacteria to switch back to an antibiotic-resistant phenotype through reactivation of MexT. This raises the possibility that cycles of antibiotic treatment are drivers for genetic diversity and adaption in the host environment. Deletion of MexT or neighbouring genomic regions has been observed in lung-adapted isolates (Warren *et al*., 2011; Smith *et al*., 2006; Hilliam *et al*., 2017) and could be a consequence of antibiotic selection in some cases. Once antibiotic resistance has been achieved through inactivation of MexS, further mutations in genes conferring antibiotic resistance, such as in *gyrA,* could arise. These subsequent mutations can work alongside increased efflux to confer very high levels of antibiotic resistance, but importantly they enable efflux to be downregulated and a switch to a more virulent phenotype to take place while the bacterium still retains an antibiotic-resistant phenotype. While it is counterintuitive that loss of a proposed detoxification system is beneficial outside of laboratory culture, in addition to increased virulence and nutrient utilisation (observed in the Mex-null PAO1 background) it may reduce acidification arising from efflux (proton import)(Olivares, Álvarez-Ortega and Martinez, 2014), repress nitrogen respiration (Arat, Bullerjahn and Laubenbacher, 2015), reduce growth arrest during the stress response (Tamman *et al*., 2015) or be compensated by redundancy in efflux systems found in *P. aeruginosa*. The fact that we do not observe reversion to an ancestral genotype where efflux is regulated by active MexS and MexT suggests that an evolutionary ratchet is operating that drives the bacteria towards switching between antibiotic-resistant and virulent phenotypes, but does not enable a return to the ancestral state, at least under laboratory conditions. The reason for this is not currently clear, but given we know that an antibiotic-sensitive disrupted MexS/MexT strain is more metabolically active than an antibiotic-resistant disrupted MexS strain, it could be hypothesised that the ancestral genotype, where MexS and MexT are fully active, retains the greatest degree of metabolic activity and therefore would be outcompeted by a disrupted MexS/MexT strain where certain metabolic pathways are not active and the bacteria do not expend resources on homeostasis. This hypothesis remains to be tested. It has been proposed that active efflux is indicative of chronic infection as the production of virulence factors that induces a host response is reduced (Lamarche and Déziel, 2011; Winstanley, O’Brien and Brockhurst, 2016) but our evidence also supports the inactivation of *mexT* as a signature of chronic infection (La Rosa, Johansen and Molin, 2018). The evolutionary trajectory may also depend upon the nature of the infection. It may be that initial activation of efflux via MexEF-OprD reduces the expression of quorum sensing-controlled virulence factors, which are subsequently kept down-regulated by mutations in *lasR,* which in turn allows inactivation of *mexT* while retaining low virulence. From a clinical point of view, knowledge of the *mexS*/*mexT* genotype of an infecting strain could be important for informing treatment. If an isolate is antibiotic-sensitive due to ancestral active MexS/MexT, then use of an antibiotic for which MexEF-OprD is a substrate could be successful but could also select for resistant mutants. However, if an isolate is antibiotic-sensitive due to inactivated MexS/MexT, then use of an antibiotic that is a MexEF-OprD substrate could potentially select for a resistant virulent strain which is less likely to be eliminated and be more damaging for the patient. As whole genome sequencing becomes more widespread within microbiological diagnostics, identifying and utilising such information will be key for guiding more targeted treatments, improving patient outcomes and preserving antibiotic efficacy.

## Methods

### Bacterial strains and culture conditions

Strains listed in Table S1 were initially grown on Columbia agar (Oxoid, UK) and then incubated at 37 °C for 16 hours at 180 rpm in supplemented M9 medium comprised of 22.2 mM glucose, 2 mM MgSO4, 0.1 mM CaCl2, 24.4 mM casamino acids and 1 mM thiamine hydrochloride (Sigma-Aldrich, USA). Strains listed in Table S3 were frozen upon receipt without laboratory passage except for Strain A which was purified on *Pseudomonas* Agar (Oxoid, UK) with cetrimide and nalidixic acid and frozen stocks were prepared after overnight growth in LB broth.

### *Generation of PA*Δ *mutant with 8-bp deletion*

DNA was extracted from PA-S (Supplementary Table S1) as per manufacturer’s instructions using the Roche MagNA pure Compact system (Roche, Switzerland), and *mexT* was PCR-amplified using Phusion DNA polymerase and Phusion GC Reaction Buffer (New England Biolabs, U.S.A.) combined with dNTPs and primers listed in Supplementary methods (Primer list A, Supplementary Methods). PCR conditions: Initial denaturation: 98 °C (5 minutes), followed by 32 cycles of: denaturation 98 °C (10 seconds), annealing 55 °C (30 seconds), elongation 72 °C (20 seconds) then final extension of 72 °C (5 minutes). Plasmid DNA from the suicide vector, pTS, was extracted using the NucleoSpin Plasmid kit (Macherey-Nagel, Germany). Amplified *mexT* DNA was digested with MfeI and BamHI and pTS DNA with EcoRI and BamHI, as per the manufacturer’s instructions (New England Biolabs Ltd, UK). The vector was treated with alkaline phosphatase and ligated with the insert at a ratio of 3:1 (insert to vector) using T4 ligase (New England Biolabs) before incubation overnight at 4 °C. The pTSmexT constructs were individually transformed into *E. coli* DH5α by heat shock: pTSmexT and *E. coli* DH5α were mixed on ice, incubated at 42 °C for 1 minute and placed on ice for 2 minutes. LB broth was added and cultures incubated at 37 °C for 2 hours. Cells were then plated onto 10 μg/mL tetracycline agar to confirm the presence of *E. coli* DH5α colonies with the construct and tetracycline resistance marker. Colony PCR (primer list A1 and A2, Supplementary Methods) and gel electrophoresis were performed to screen colonies for the required insert. PCR steps included: 95 °C for 5 minutes, 30 cycles of 95 °C for 30 seconds, 55 °C for 30 seconds, 72 °C for 2 minutes and then 72 °C for 10 minutes. Electrocompetent PA cells were prepared by washing in 300 mM sucrose and transformation with the constructs was performed using a Gene Pulser Electroporation System (Bio-Rad, U.S.A.) with the following settings: 200 Ω, 2.5 kV before plating onto sucrose agar to counter select the plasmid. Colonies with the insert were then confirmed by sequencing (Eurofins Scientific, Luxembourg) and thereafter named PAΔ.

### Generation of PAn

To generate PAn, PA overnight cultures were diluted to 10^6^ CFU/mL and plated onto LB agar (Oxoid, UK) containing 0.25 µg/mL ciprofloxacin. One colony, termed PAn, was identified by PCR (*Primer list B, Supplementary Methods)* with a single X repeat sequence. PCR conditions included: 95 °C for 10 sec, 60 °C for 30 sec (45 cycles) followed by a melt curve analysis. The single X repeat sequence was also confirmed using whole genome sequencing.

### Motility testing

Swarming, swimming and twitching phenotypes were tested on LB agar concentrations of 0.3 %, 0.5 % and 1 % respectively (O’May and Tufenkji, 2011; Rashid and Kornberg, 2000). For swarm plates, 5 μL of the inoculum was placed onto the agar surface. For swimming tests, 5 μL of inoculum representing 10^8^ CFU/mL, was stabbed into the centre of the agar. For twitching, the inoculum was pelleted and a toothpick used to inoculate the agar-petri dish interface. Plates were incubated for 18 h at 37 °C before the diameters of the motility zones were measured.

### Antibiotic minimal inhibitory concentration (MIC) testing

Ciprofloxacin stocks were prepared in 0.1 M HCl, chloramphenicol stocks were prepared in ethanol and imipenem stocks were prepared in MOPs buffer. MICs for *P. aeruginosa* strains against imipenem, ciprofloxacin and chloramphenicol (Sigma, UK) were determined by an agar incorporation method. Isolates were streaked onto Mueller-Hinton agar and incubated overnight at 37 °C. A single colony was used to inoculate Mueller-Hinton broth and cultured overnight at 37 °C. Overnight cultures were normalised to an OD_600_ of 0.5 MacFarland standard and 1 µL inoculum stamped onto Mueller-Hinton plates containing imipenem (range 0.125 – 128 µg/mL), ciprofloxacin (range 0.125 – 128 µg/mL) or chloramphenicol (range 1 – 2048 µg/mL) alongside an antibiotic-free growth control plate using a multi-point inoculator (Denley, UK). MICs were read as the concentrations that showed no visible growth or haze after overnight incubation at 37 °C. Each strain was tested in duplicate. *P. aeruginosa* ATCC 27853 and *E. coli* ATCC 25922 were included as control strains.

### Phenotype microarray

Utilisation of 626 substrates was tested using phenotype microarray (PM) plates and protocols supplied by BioLog Inc, USA. Briefly, strains were serially cultured on Columbia agar twice with incubation overnight at 37 °C. Bacterial colonies were suspended in inoculation fluid-0 and dye A, and 100 µL was aliquoted into each well of PM plates 1-2. Sodium succinate (0.54 g/mL) and ferric citrate (0.049 mg/mL) were added to the inoculating fluid for PM plates 3-8. Plates were incubated at 30 °C for 96 hours in the OmniLog reader. The signal value (SV) (Homann *et al*., 2005) for each substrate was calculated and negative controls subtracted from the results. Resultant negative values were assigned a value of 0, indicating no growth. To enable fold change (FC) calculations, all results (negative and positive) were adjusted by adding a value of 1. SVs for each substrate were averaged across replicates for each strain and the FC calculated between PA versus PA*Δ* and PA versus PAn. A normal distribution was applied. Results outside of the 95 % confidence interval were subjected to a paired Student’s T-test and those with a P value of < 0.05 were considered significant.

### Analysing growth characteristics

Strains reverting to a reduced chloramphenicol tolerance were initially cultured overnight in the media the reversion was generated in (BHI for Strain 10; revertants A and B, Strain 10; revertant C and D, SSM9PR for the revertant of Strain A) at 37 °C alongside its respective chloramphenicol sensitive parent and chloramphenicol tolerant mutant. The density of the overnight culture was adjusted to OD_600_ 0.08-0.1 and further diluted 1:20 to create an inoculum containing approximately 5×10^6^ cfu/mL. 20 µL of the final inoculum was then added to five wells of a 96 well plate containing 180 µL of broth and five wells containing 180 µL of broth supplemented with a concentration of chloramphenicol two doubling dilution below the lowest MIC detected for each parent-mutant-revertant strain set (8 µg/mL for both BHI and SSM9PR). Only the inner wells of the 96 well plate were used to avoid condensation interfering with the readings. The plate was incubated at 37 °C with shaking at 200 rpm at the beginning of each cycle on the BMG FLUOstar® Omega plate reader. Absorbance readings at a wavelength of 600 nm were taken every 5 minutes over the course of 24 hours. Growthcurver (Sprouffske and Wagner, 2016) on R v4.0.0 was used to calculate the area under the experimental curve (*auc*). Mann-Whitney *U* tests were used to compare the means and P-values were adjusted for multiple testing by the Benjamini-Hochberg method with Rstatix v0.7.0 (Kassambara). Growth curves were plotted with ggplot2 v3.3.5 (Wickham, 2009) using the average OD_600_ at 15 minute intervals up to 16.5 hours to cover the exponential phase of growth. Error bars are shown as 95 % confidence intervals. The experiment was conducted twice with 5 technical replicates included each time.

### Virulence assays

The relative virulence of the *P. aeruginosa* strains PA, PAn and PAΔ was assessed in the *Galleria mellonella* model according to a protocol modified from McMillan et al (2015) (McMillan *et al*., 2015). Briefly, larvae (UK Waxworms Ltd, UK) in groups of 12 were injected with PBS suspensions containing ca. 10 total CFU. In addition, one control group underwent no manipulation to control for background larval mortality (no manipulation control) while another group (uninfected control) was injected with PBS only to control for the impact of physical trauma. Larvae were kept in petri dishes in the dark at 37 °C for up to 24 hours and inspected every 6 hours so that percentage survival could be calculated for each group. Larvae were considered dead if they did not move after being stimulated with a sterile inoculation loop. The experiment was repeated in triplicate.

### DNA extraction and sequencing

For PAO1-PA, PAn and PAΔ, per strain, DNA was extracted from 1 mL of overnight supplemented M9 culture using the MagNA pure Bacterial lysis kit with RNAse (Qiagen, Germany) on the MagNA Pure Compact instrument (Roche, Switzerland). 1 ng of DNA was used in a standard Nextera XT library preparation for paired-end sequencing on the MiSeq with v3 chemistry (Illumina, U.S.A). For DNA extraction from chloramphenicol-resistant mutants, revertants and the corresponding parent, DNA was extracted from 1 mL of overnight LB culture (supplemented with 100 µg/mL chloramphenicol for chloramphenicol-resistant mutants) using the GeneJet Genomic DNA Purification (ThermoFisher, UK). 1 ng of DNA was used in a standard Nextera XT library preparation before sequencing on the NextSeq with v2 chemistry (Illumina, U.S.A). Adapter and quality trimming of reads was performed with Trimmomatic (Bolger, Lohse and Usadel, 2014).

### DNA and protein analysis

To identify single nucleotide variants in PAO1 isolates, DNA sequence reads in fastq format were aligned to the reference genome - *P. aeruginosa* PAO1, obtained from NCBI Genbank with accession number NC_002516. For non-PAO1 isolates used in the evolution experiments, SNP calls for mutants and revertants was based on alignment to their parent genomes. Reads were aligned to the reference using Bowtie2 (Langmead and Salzberg, 2012) and alignments were sorted, indexed and stored in bam file format using samtools (Li *et al*., 2009). Variants were called using Bayesian inference with freebayes v1.3.1 (Garrison and Marth, 2012). Variants were also called with samtools variant calling methods. High confidence changes had a samtools quality score of >200. All other changes were low confidence, with quality scores of <45 and were excluded. Freebayes confirmed the high confidence changes. Spurious variants were excluded by calling variants of self reads against parent reference, these were intersected using bcftools and intersecting variants were removed from the mutant and revertant variant calls. A further step of filtering variants with vcfutils varFilter was carried out. SNPEff v4.3t was subsequently run across the filtered vcf files and results aggregated into a table (Cingolani *et al*., 2012).

The ARIBA tool (Hunt *et al*., 2017) was used to identify changes in specified genes within chloramphenicol-resistant mutants and chloramphenicol-sensitive revertants with *mexS* and *mexT* (and *cmrA*) from *P. aeruginosa* UCBPP-PA14 (obtained from the Pseudomonas Genome Database (Winsor *et al*., 2016)) as query genes. As a further check, protein sequences of MexS and MexT were extracted from each strain using BLAST with UCBPP-PA14 sequences as query and aligned.

Other analyses of genome sequences, obtained from the National Center for Biotechnology Information (NCBI) and *Pseudomonas* Genome Database, were performed using the NCBI BLAST and Clustal Omega (Sievers *et al*., 2011) tools. In addition, a Panaroo (Tonkin-Hill *et al*., 2020) analysis was carried out with reference strains PAO1, PA14, LES together with the mutant and revertant sequences.

### COG mapping

COG annotations were downloaded from the *Pseudomonas* Genome Database (last accessed November 2017) and used to categorise the differentially expressed genes in PAΔ and PAn. Some genes have not yet been annotated with COG categories: this included 47 and 132 genes up and down-regulated in PAΔ and 43 and 111 genes up and down-regulated in PAn.

### RNA extraction

Five colonies from each strain grown on Columbia agar were used to inoculate 10 mL of Lysogeny Broth (LB) and incubated at 37 °C for 24 hours. Cultures were diluted to 1:1000 in fresh M9 and grown for 24 hours at 37 °C, 180 rpm then diluted 1:100 in fresh M9 and incubated for 5 hours to ensure log phase. Cultures were then mixed with RNAprotect Bacterial Reagent and centrifuged according to the manufacturer’s protocol (Qiagen, Netherlands). Bacterial pellets were stored at –80°C. TE buffer (10 mM Tris, 1 mM EDTA, pH 8.0) containing 15 mg/mL lysozyme (Fisher Scientific Ltd, UK) and proteinase K (Roche, Switzerland) were added to cell pellets and incubated for 10 minutes at ambient temperature. Samples were then processed using the RNeasy mini Kit (Qiagen) according to the manufacturer’s protocol, with the inclusion of on-column DNAse treatment from the RNase free DNase kit (Qiagen). Samples were also treated with DNase using the Turbo DNA-free Kit (Life Technologies Ltd, UK) and washed with the RNeasy mini kit. Three independent samples were processed per strain.

### RNA sequencing

Samples were sequenced by Genomed, Poland. Ribo-depletion was performed using the Ribo-Zero rRNA removal kit (Bacteria) (Epicentre, U.S.A) and libraries prepared using the NEBNext® Ultra™ RNA Library Preparation Kit. Sequencing was carried out using Hi-Seq (Illumina, U.S.A) with paired-end sequencing (2× 100 bp) and V3 chemistry reagents.

### RNA sequencing analysis

Transcriptome reads were aligned to the reference genome (NCBI Genbank with accession number NC_002516) by Genomed using TopHat (Kim *et al*., 2013) and stored in samtools bam file format. Results were provided in FPKM value output. Reads were counted and normalised in R using simple log fold change, edgeR (Robinson, McCarthy and Smyth, 2010) glm, and edgeR classic formats, as well as using DESeq2 (Love, Huber and Anders, 2014) methods. Results from the analyses were output as sorted Excel files. DESeq2 output was used for downstream analysis after verification that the other transcriptomics methods provided similar results. P-values of 0.05 or less were calculated by applying a Student’s t-test and a Bonferroni correction. To account for mapping all data to the reference strain (PAO1 reference in the Pseudomonas Genome Database (Winsor *et al*., 2016), in the XXY formation), *mexT* was split into two regions for this analysis. The first region was from the gene start (position 2807469) and included the XXY repeat - the second region continued on from this to the end of the gene. Results were ranked according to log_2_ fold change differences between PA and PAΔ or PA and PAn. Transcriptome assemblies were generated using Trinity (Haas *et al*., 2013).

### Transcriptional regulatory network reconstruction

DNA binding sites for *P. aeruginosa* PAO1 transcription factors (TF) were retrieved from KEGG. For any given TF, we inferred Position Specific Scoring Matrices (PSSMs) representing conserved binding motifs from the literature-sourced raw binding sites using the *consensus* tool (Thomas-Chollier *et al*., 2011; Thomas-Chollier *et al*., 2008). PSSMs were then converted to *transfac* format using the *convert-matrix* tool (Kiliç *et al*., 2014; Münch *et al*., 2003). To check if any TFs had an effect on gene expression from binding to intergenic regions, we took 2000 bp upstream from the start codon of every gene in the PAO1 reference genome, with no overlap to the upstream ORFs (operon information was obtained from operonDB (Pertea *et al*., 2009). The *transfac*-formatted PSSMs were used to scan upstream sequences with the *matrix-scan* tool. Putative hits (i.e. genes or operons harbouring TF binding sites) with a p-value < 1E-05 were considered significant.

### Selection of spontaneous chloramphenicol-resistant P. aeruginosa

Clinical *P. aeruginosa* isolates were streaked on Tryptic Soy Agar (TSA) and incubated overnight at 37 °C. Five colonies were resuspended in 2 mL PBS and 10 µL spotted onto TSA and TSA supplemented with chloramphenicol at 100, 200 or 400 µg/mL before overnight incubation at 37 °C. In parallel, 25µL of un-passaged clinical *P. aeruginosa* stocks were spread directly onto TSA plates supplemented with 400 µg/mL chloramphenicol alongside a TSA growth control and incubated at room temperature for > 48 hours. Putative chloramphenicol resistant mutants were streaked on TSA supplemented with chloramphenicol at 400 µg/mL and incubated at 37 °C alongside the parent (Table S3). Confirmed resistant mutants were harvested from plates in 0.5 mL LB and supplemented with 25 % aq. glycerol for storage at -80 °C.

### Selection of antibiotic-sensitive revertants

Chloramphenicol-resistant *P. aeruginosa* were streaked on TSA supplemented with chloramphenicol at 400 µg/mL and approximately 5 colonies used to inoculate either 1 mL Brain Heart Infusion (BHI) or SSM9PR (1× M9 salts, 2 mM MgSO4, 0.1 mM CaCl2, 1 % glucose, 1 % casamino acids, 1 mM thiamine HCl, 0.05 mM nicotinic acid). The *P. aeruginosa* PA, PAn and PAΔ strains were included as growth controls. Cultures were passaged up to 6 times over 2.5 weeks with incubation at either room temperature (static) or 37 °C (180 rpm) in the intervening time. For passage, 10 µL culture was used to inoculate fresh BHI or SSM9PR media in the same vial. At 3, 5 and 6 days, 2 µL culture was replica plated on chloramphenicol (400 µg/mL) and antibiotic-free plates (alongside the parent) and isolates with visibly reduced growth on chloramphenicol were streaked on TSA for single colonies and incubated at 37 °C overnight. Single colonies (4-10 cfu) from streaked cultures were replica plated on chloramphenicol (400 µg/mL) and antibiotic-free media to identify susceptible revertants which were confirmed by MIC testing. Putative revertants were harvested from TSA plates in 0.5 mL LB and supplemented with 25 % glycerol and stored at -80 °C.

## Data Availability

All Illumina data have been deposited at the European Nucleotide Archive (www.ebi.ac.uk/ena) under project PRJEB18543 (ERP020481) for the transcriptomics data and PRJEB28534 (ERP110744) for the mutants and revertants sequencing data.

## Acknowledgements

We thank David Livermore for useful discussions about the concept of the study. AC was supported by a PhD studentship from the Norwich Medical School of the University of East Anglia. TK and PS were supported by a fellowship to TK in computational biology at the Earlham Institute (BB/CSP1720/1) in partnership with the Quadram Institute Bioscience, and strategically supported by the Biotechnology and Biological Sciences Research Council, UK (BB/J004529/1; BB/R012490/1 and its constituent projects BBS/E/F/000PR10353 and BBS/E/F/000PR10355). SA and MU were supported by the Erasmus programme. JGM was supported by Institute Strategic Program Grant BB/X010996/1 to the John Innes Centre. Research presented in this paper was in part carried out on the High Performance Computing Cluster supported by the Research and Specialist Computing Support service at the University of East Anglia. The later stages of this work were supported by funding from the Norwich Medical School of the University of East Anglia.

We thank the microbiology department at the Norfolk and Norwich University Hospital for provision of *P. aeruginosa* isolates and associated resistance data supplied through the Norwich Biorepository.

## Supplementary Datasets

**Dataset S1. RNA-Seq analysis of P. aeruginosa strains**

RNA-Seq was carried out by an external company which provided the count data. Count data was further analysed using the R program DESeq2 to provide log2 fold change results and *p*-values on the PAO1 backbone sequence. The resultant count dataset is provided in the table. On manual examination of RNA-seq read alignments on the *mexT* gene, it could be seen that the N-terminal and C-terminal portions of the gene had differing numbers of reads. To examine this further in the RNA-seq data, the gene was split into two parts, here denoted 1PA2492 for the N-terminal part and 2PA2492 for the C-terminal part.

**Dataset S2. Differential gene expression analysis and COG classification in P. aeruginosa strains**

This dataset presents differential gene expression analyses between PA, PAΔ, and PAn, based on reads aligned to the PAO1 reference genome (NC_002516). Transcript abundance was quantified using DESeq2, and genes with log2FC > -1 and < 1 were excluded. Differential expression was assessed using adjusted *p*-values (*p* ≤ 0.05, Bonferroni-corrected). The dataset includes the following worksheets: Shared Down-Regulated Genes: Genes down-regulated in both PAΔ and PAn vs. PA. Shared Up-Regulated Genes: Genes up-regulated in both PAΔ and PAn vs. PA. PAΔ vs. PA: Genes differentially expressed in PAΔ vs. PA. PAn vs. PA: Genes differentially expressed in PAn vs. PA. COG Classification: Summary of differentially expressed genes by COGs.

**Dataset S3. Transcriptional Regulatory Network of P. aeruginosa PAO1.**

Regulatory network of *P. aeruginosa* PAO1, reconstructed using transcription factor binding site data from KEGG and literature-derived PSSMs. Upstream intergenic regions (2000 bp) were scanned for putative TF binding sites using matrix-scan, with significant hits identified at *p* < 1E-05. Data is presented in each worksheet: Target Lists: Genes predicted to be regulated by MexT, CifR, and PvdS. MexT Targets-Operons : MexT-regulated operons, with binding sites marked in red. CifR Targets-Operons: CifR-regulated operons, with binding sites marked in red. PvdS Targets-Operons: PvdS-regulated operons, with binding sites marked in red. COG Classification of Targets: Functional categorization of MexT, CifR, and PvdS targets based on COGs.

**Dataset S4. Fold change of substrates differentially utilised by the mexT variants (* p <0.05), n=2**

Fold change and utilization of substrates significantly affected by *mexT* variants (*p* < 0.05, n = 2), assessed using phenotype microarray (PM) analysis. Growth on 626 substrates was measured over 96 hours, and signal values were normalized and analysed. Fold changes were calculated for PA vs. PAΔ and PA vs. PAn, with significance determined at *p* < 0.05. Significant Results: Substrates showing significant differential utilization among *mexT* variants. All Results: Raw and processed SV data for all tested substrates, including FC values and *p*-values.

**Dataset S5. SNP analysis of P. aeruginosa strains, chloramphenicol-resistant mutants and passaged revertants/non-revertants**

SNP analysis was carried out by calling variants on read alignments with Freebayes and Samtools on genomes annotated with Panaroo. The program SNPEff was used to analyse variant effects into SNPEff defined categories with impacts HIGH, LOW, MODERATE and MODIFIER. These impacts are described in detail in the SNPEff documentation. In accordance with the SNPEff outputs, these impacts may not be mutually exclusive.

**Figure S5.** Heatmap of the the data from dataset S5 focussed on the *mex* operon. The heatmap was formed from the dataset readout using R libraries to aggregate the data according to SNPEff variant type and provide total counts.

