## Supplementary Information for "Plasticity in a bacterial global regulatory switch that drives a shift in antibiotic resistance and virulence"

**Methods**

**Primer list A1: Primers used to construct PAΔ**

mexT forward: ATGGATCCGTTCGAAGCCGAGACCG

mexT reverse: ATGAATTCCTCCTCGTCGACGAAGC

**Primer list A2: Primers used to construct PAΔ**

pTS forward: CGGCAGGTATATGTGATGGG

pTS reverse: CCATGAGTGACGACTGAATCCG

**Primer list B: Primers used to study the presence of a single X repeat sequence in PAn**

**Forward:** CGCAGAGAAACTGTTCCT

**Reverse:** GGTACGGACGAACAGC

**Supplementary tables**

**Supplementary Table S1. *P. aeruginosa* PAO1 strains and plasmids used in this study**

| Strain/Plasmid | Genotype or relevant characteristic | Source |
| --- | --- | --- |
| PA | PA parent strain with two X repeats in *mexT* | John Innes Centre |
| PAΔ | PA engineered with one X repeat in *mexT* | This study |
| PAn | PA with one X repeat in *mexT* selected for on sub-MIC ciprofloxacin agar (nfxC-phenotype) | This study |
| PA-S | PAO1 with an 8-bp deletion in the *mexT* gene. Used in allelic exchange to genetically engineer PAΔ. | Derived from ATCC 15692 |
| pTS | 8.8-kb broad-host-range cloning/shuttle vector (Tet^r^*,oriColE1, sacB)* | Dr Jacob Malone, John Innes Centre |

**Supplemental Table S2. Survey of *mexT* sequences in *P. aeruginosa***

| **Source** | **Isolate** | ***mexT* genotype** | **Strain origin** | **Reference(s)** |
| --- | --- | --- | --- | --- |
| Literature | mPAO1/P1, L-PAO1, C-PAO1, Ig-PAO1, Ma-PAO1, Geneva, PAO1-UT | XY | Wound isolate,  laboratory reference strain | Holloway, 1955;  Maseda *et al*., 2000;  Stover *et al*., 2000; Klockgether *et al*., 2010, Luong *et al*., 2014; Sidorenko *et al*., 2017 |
|  | PAO1-UW (Reference), R-PAO1, PAO1-L, PAO1S-Lac, PAO1-GP | XXY |  |  |
|  | mPAO1/P2, H-PAO1, PAO1-DSM (1707), PAO1-Tokai | XY* |  |  |
|  | CF non-mutator isolate (n = 1) | Deletion | Cystic fibrosis isolate | Warren *et al*., 2011 |
|  | Cystic fibrosis isolate | XY*/  Deletion | Cystic fibrosis isolate | Smith *et al*., 2006 |
|  | Bronchiectasis (n = 4) | Deletion | Bronchiectasis isolate | Hilliam *et al*., 2017 |
| NCBI database (*mexT* XXY) | C7447m | XXY | Mucoidy cystic fibrosis isolate, Vancouver, BC | Yin *et al*., 2013 |
|  | PA1R | XXY | Phage-resistant derivative of the PA1 (XY) respiratory isolate | Le *et al*., 2014 |
|  | *Pseudomonas aeruginosa* M10 | XXY | Isolated from a screen for biosurfactant producing bacteria at the Churince water spring, Mexico | [ATAG01000457.1](https://www.ncbi.nlm.nih.gov/nuccore/ATAG01000457) |
|  | *Pseudomonas aeruginosa* B3-1811 | XXY | Cystic fibrosis patient undergoing chemotherapy/ Denmark | [CBMP010000089.1](https://www.ncbi.nlm.nih.gov/nuccore/CBMP010000089) |

**Supplemental Table S3. *P. aeruginosa* strain collection**

| **Sequencing sample accession** | **Strain name** | **Details** |
| --- | --- | --- |
| SAMEA6618357 | Strain 1 | Clinical ear swab |
| SAMEA6618338 | Strain 1 Cm^R^ | *In vitro* chloramphenicol-resistant mutant of Strain 1 |
| SAMEA6618311 | Strain 1 Revertant (BHI) | Chloramphenicol-sensitive revertant of Strain 1 Cm^R^ after BHI passage |
| SAMEA6618318 | Strain 1 Non-revertant (BHI) | Chloramphenicol-resistant non-revertant of Strain 1 Cm^R^ after BHI passage |
| SAMEA6618321 | Strain 1 Non-revertant A (M9) | Chloramphenicol-resistant non-revertant of Strain 1 Cm^R^ after M9 passage |
| SAMEA6618308 | Strain 1 Non-revertant B (M9) | Chloramphenicol-resistant non-revertant of Strain 1 Cm^R^ after M9 passage |
| SAMEA6618352 | Strain 10 | Clinical mid stream urine isolate |
| SAMEA6618347 | Strain 10 Cm^R^ 1 | *In vitro* chloramphenicol-resistant mutant of Strain 10 |
| SAMEA6618333 | Strain 10 Revertant A (BHI) | Chloramphenicol-sensitive revertant of Strain 10 Cm^R^1 after BHI passage |
| SAMEA6618334 | Strain 10 Revertant B (BHI) | Chloramphenicol-sensitive revertant of Strain 10 Cm^R^1 after BHI passage |
| SAMEA6618343 | Strain 10 Cm^R^ 2 | *In vitro* chloramphenicol-resistant mutant of Strain 10 |
| SAMEA6618329 | Strain 10 Revertant C (BHI) | Chloramphenicol-sensitive revertant of Strain 10 Cm^R^2 after BHI passage |
| SAMEA6618330 | Strain 10 Revertant D (BHI) | Chloramphenicol-sensitive revertant of Strain 10 Cm^R^2 after BHI passage |
| SAMEA6618353 | Strain 11 | Clinical catheter urine |
| SAMEA6618344 | Strain 11 Cm^R^ | *In vitro* chloramphenicol-resistant mutant of Strain 11 |
| SAMEA6618331 | Strain 11 Revertant A (BHI) | Chloramphenicol-sensitive revertant of Strain 11 Cm^R^ after BHI passage |
| SAMEA6618332 | Strain 11 Revertant B (BHI) | Chloramphenicol-sensitive revertant of Strain 11 Cm^R^ after BHI passage |
| SAMEA6618358 | Strain 20 | Clinical ear swab |
| SAMEA6618349 | Strain 20 Cm^R^ | *In vitro* chloramphenicol-resistant mutant of Strain 20 |
| SAMEA6618323 | Strain 20 Revertant A (BHI) | Chloramphenicol-sensitive revertant of Strain 20 Cm^R^ after BHI passage |
| SAMEA6618324 | Strain 20 Revertant B (BHI) | Chloramphenicol-sensitive revertant of Strain 20 Cm^R^ after BHI passage |
| SAMEA6618307 | Strain A | Clinical isolate from blood |
| SAMEA6618339 | Strain A Cm^R^ | *In vitro* chloramphenicol-resistant mutant of Strain A |
| SAMEA6618309 | Strain A Revertant (M9) | Chloramphenicol-sensitive revertant of Strain A Cm^R^ after M9 passage |
| SAMEA6618310 | Strain A Non-revertant (M9) | Chloramphenicol-resistant non-revertant of Strain A Cm^R^ after M9 passage |
| SAMEA6618319 | Strain 1 passage A (BHI) | Control: Strain 1 parent passaged in BHI |
| SAMEA6618320 | Strain 1 passage B (BHI) | Control: Strain 1 parent passaged in BHI |
| SAMEA6618312 | Strain 1 passage A (M9) | Control: Strain 1 parent passaged in M9 |
| SAMEA6618313 | Strain 1 passage B (M9) | Control: Strain 1 parent passaged in M9 |
| SAMEA6618361 | Strain 4 | Clinical catheter urine |

**Supplementary figures**

**Supplemental Figure S1: Phenotypic characterisation of parent and *mexT* variant strains**

**A. Swarm, B. Swim, and C. Twitch zone diameters measured for the parent and *mexT* variant strains (n=3, error bars = sd). D. Kaplan-Meier plot of *Galleria mellonella* survival during 24 h at 37°C after injection with PA, PAn and PAΔ.** The accurate inoculum doses were 7.27 ±0.23 CFU (mean ± standard error) for PA, 7.91 ±1.80 CFU for PAn and 7.42 ±1.96 CFU for PAΔ. Experiments were performed with three independent batches of larvae and a group size of 12 per replicate. Two-tailed logrank tests confirmed a significant difference in virulence between PA and PAΔ (*p*=3.27e-7) and PAn and PAΔ (*p*=4.02e-4), but not between PA and PAn (*p*=0.077). For clarity, data are not shown for groups of non-injected larvae (no manipulation control) or injected with phosphate-buffered saline carrier solvent only. *n*=36.

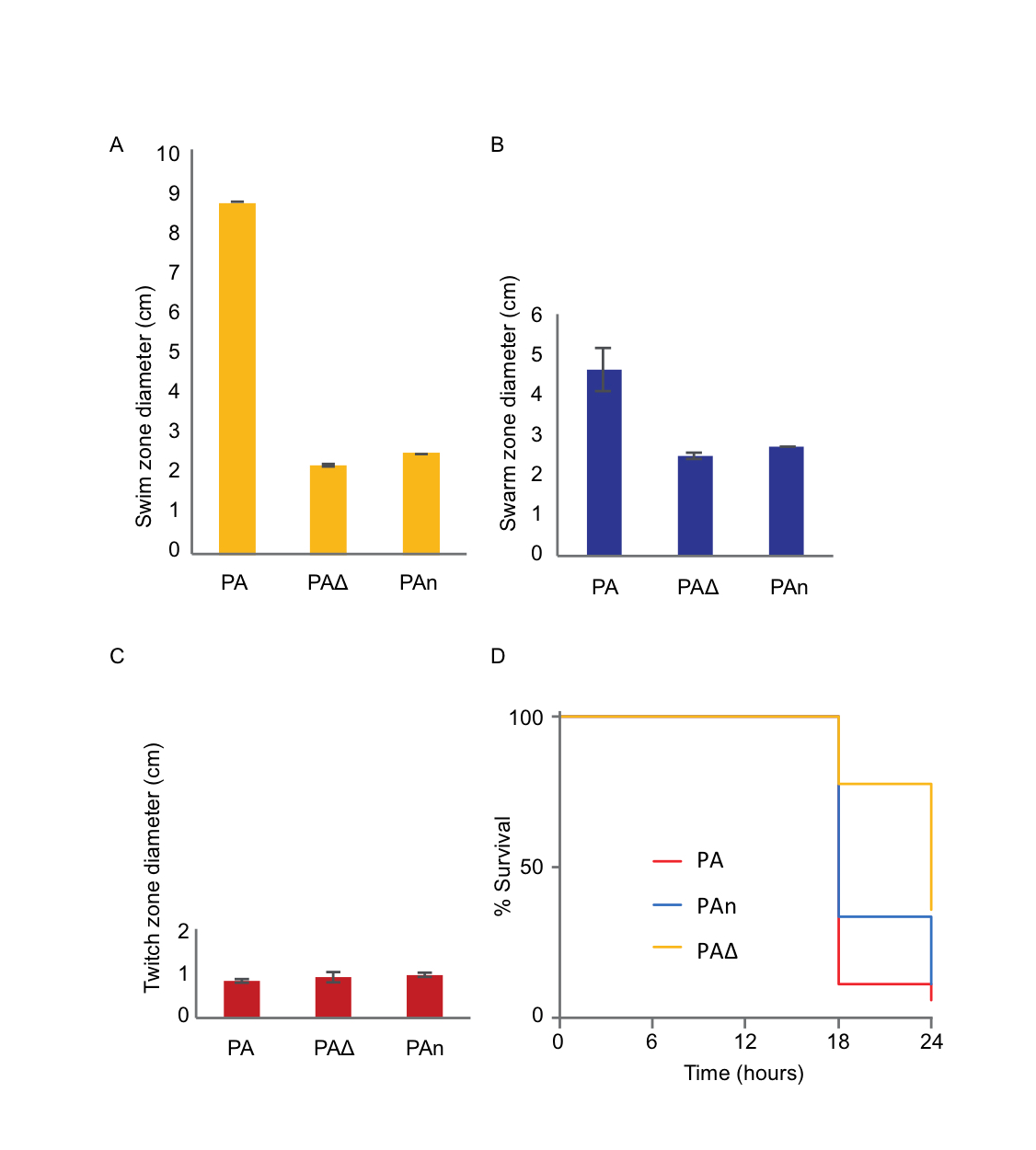

**Supplemental Figure S2: Multiple alignment of MexS sequences from PAO1 lineages and relevant *P. aeruginosa* isolates.** An inactivating N249D mutation (highlighted) is present in MexS in most PAO1 lineages (represented here by PAO1-UW) but absent from three reported PAO1 lineages (PAO1-VE2, PAO1-VE13 and PAO1H2O). The mutation is not present in the *P. aeruginosa* c7447m and PA1R strains which are *mexT* XXY type or in the non-PAO1 strains PA14 and LESB58 which were included for comparison. Multiple alignment was performed using Clustal Omega (EMBL-EBI, Sievers *at el*., 2011). Sequences were obtained from the *Pseudomonas* database (February 2020, Winsor *et al*., 2016): for cases where MexS was not annotated, the PAO1 (reference) MexS protein sequence was used as input in a BLAST search to extract the correct protein.

```
CLUSTAL O(1.2.4) multiple sequence alignment

PAO1_Reference_GCF_000006765.1|latest_PA2491_mexS      MSRVIRFHQFGPPEVLKCEELPTPAPAAGEVLVRVQAIGVSWKDVLWRQNLAPEQAALPS	60
LESB58_GCF_000026645.1|latest_PALES_28051              MSRVIRFHQFGPPEVLKCEELPTPAPAAGEVLVRVQAIGVSWKDVLWRQNLAPEQAALPS	60
PAO1H2O_GCF_00071515.1|latest_OU9_02515                MSRVIRFHQFGPPEVLKCEELPTPAPAAGEVLVRVQAIGVSWKDVLWRQNLAPEQAALPS	60
PAO1-VE2_GCF_000484495.2|latest_N296_2562              MSRVIRFHQFGPPEVLKCEELPTPAPAAGEVLVRVQAIGVSWKDVLWRQNLAPEQAALPS	60
PAO1-VE13_GCF_000484545.2|latest_N297_2562             MSRVIRFHQFGPPEVLKCEELPTPAPAAGEVLVRVQAIGVSWKDVLWRQNLAPEQAALPS	60
UCBPP-PA14_GCF_000014625.1|latest_PA14_32420           MSRVIRFHQFGPPEVLKCEELPTPAPAAGEVLVRVQAIGVSWKDVLWRQNLAPEQAALPS	60
C7447m_GCF_000468935.2|latest_M802_2559                MSRVIRFHQFGPPEVLKCEELPTPAPAAGEVLVRVQAIGVSWKDVLWRQNLAPEQAALPS	60
PA1_GCF_000496605.2|latest_PA1S_RS13175                MSRVIRFHQFGPPEVLKCEELPTPAPAAGEVLVRVQAIGVSWKDVLWRQNLAPEQAALPS	60
PA1R_GCF_000496645.1|latest_PA1R_gp0285                MSRVIRFHQFGPPEVLKCEELPTPAPAAGEVLVRVQAIGVSWKDVLWRQNLAPEQAALPS	60
                                                       ************************************************************

PAO1_Reference_GCF_000006765.1|latest_PA2491_mexS      GLGFELAGEVLAVGAGVGDLPLGSRVASFPAHTPDHYPAYGDVVLMPRAALAVYPEVLTP	120
LESB58_GCF_000026645.1|latest_PALES_28051              GLGFELAGEVLAVGAGVGDLPLGSRVASFPAHTPDHYPAYGDVVLMPRAALAVYPEVLTP	120
PAO1H2O_GCF_00071515.1|latest_OU9_02515                GLGFELAGEVLAVGAGVGDLPLGSRVASFPAHTPDHYPAYGDVVLMPRAALAVYPEVLTP	120
PAO1-VE2_GCF_000484495.2|latest_N296_2562              GLGFELAGEVLAVGAGVGDLPLGSRVASFPAHTPDHYPAYGDVVLMPRAALAVYPEVLTP	120
PAO1-VE13_GCF_000484545.2|latest_N297_2562             GLGFELAGEVLAVGAGVGDLPLGSRVASFPAHTPDHYPAYGDVVLMPRAALAVYPEVLTP	120
UCBPP-PA14_GCF_000014625.1|latest_PA14_32420           GLGFELAGEVLAVGAGVGDLPLGSRVASFPAHTPDHYPAYGDVVLMPRAALAVYPEVLTP	120
C7447m_GCF_000468935.2|latest_M802_2559                GLGFELAGEVLAVGAGVGDLPLGSRVASFPAHTPDHYPAYGDVVLMPRAALAVYPEVLTP	120
PA1_GCF_000496605.2|latest_PA1S_RS13175                GLGFELAGEVLAVGAGVGDLPLGSRVASFPAHTPDHYPAYGDVVLMPRAALAVYPEVLTP	120
PA1R_GCF_000496645.1|latest_PA1R_gp0285                GLGFELAGEVLAVGAGVGDLPLGSRVASFPAHTPDHYPAYGDVVLMPRAALAVYPEVLTP	120
                                                       ************************************************************

PAO1_Reference_GCF_000006765.1|latest_PA2491_mexS      VEASVYYTGLLVAYFGLVDLAGLKAGQTVLITEAARMYGPVSIQLAKALGARVIASTKSA	180
LESB58_GCF_000026645.1|latest_PALES_28051              VEASVYYTGLLVAYFGLVDLAGLKAGQTVLITEAARMYGPVSIQLAKALGARVIASTKSA	180
PAO1H2O_GCF_00071515.1|latest_OU9_02515                VEASVYYTGLLVAYFGLVDLAGLKAGQTVLITEAARMYGPVSIQLAKALGARVIASTKSA	180
PAO1-VE2_GCF_000484495.2|latest_N296_2562              VEASVYYTGLLVAYFGLVDLAGLKAGQTVLITEAARMYGPVSIQLAKALGARVIASTKSA	180
PAO1-VE13_GCF_000484545.2|latest_N297_2562             VEASVYYTGLLVAYFGLVDLAGLKAGQTVLITEAARMYGPVSIQLAKALGARVIASTKSA	180
UCBPP-PA14_GCF_000014625.1|latest_PA14_32420           VEASVYYTGLLVAYFGLVDLAGLKAGQTVLITEAARMYGPVSIQLAKALGARVIASTKSA	180
C7447m_GCF_000468935.2|latest_M802_2559                VEASVYYTGLLVAYFGLVDLAGLKAGQTVLITEAARMYGPVSIQLAKALGARVIASTKSA	180
PA1_GCF_000496605.2|latest_PA1S_RS13175                VEASVYYTGLLVAYFGLVDLAGLKAGQTVLITEAARMYGPVSIQLAKALGARVIASTKSA	180
PA1R_GCF_000496645.1|latest_PA1R_gp0285                VEASVYYTGLLVAYFGLVDLAGLKAGQTVLITEAARMYGPVSIQLAKALGARVIASTKSA	180
                                                       ************************************************************

PAO1_Reference_GCF_000006765.1|latest_PA2491_mexS      EEREFLREQGADKVVVTDEQDLVLEVERFTEGKGVNVILDELGGPQMTLLGDVSATRGKL	240
LESB58_GCF_000026645.1|latest_PALES_28051              EEREFLREQGADKVVVTDEQDLVLEVERFTEGKGVNVILDELGGPQMTLLGDVSATRGKL	240
PAO1H2O_GCF_00071515.1|latest_OU9_02515                EEREFLREQGADKVVVTDEQDLVLEVERFTEGKGVNVILDELGGPQMTLLGDVSATRGKL	240
PAO1-VE2_GCF_000484495.2|latest_N296_2562              EEREFLREQGADKVVVTDEQDLVLEVERFTEGKGVNVILDELGGPQMTLLGDVSATRGKL	240
PAO1-VE13_GCF_000484545.2|latest_N297_2562             EEREFLREQGADKVVVTDEQDLVLEVERFTEGKGVNVILDELGGPQMTLLGDVSATRGKL	240
UCBPP-PA14_GCF_000014625.1|latest_PA14_32420           EEREFLREQGADKVVVTDEQDLVLEVERFTEGKGVNVILDELGGPQMTLLGDVSATRGKL	240
C7447m_GCF_000468935.2|latest_M802_2559                EEREFLREQGADKVVVTDEQDLVLEVERFTEGKGVNVILDELGGPQMTLLGDVSATRGKL	240
PA1_GCF_000496605.2|latest_PA1S_RS13175                EEREFLREQGADKVVVTDEQDLVLEVERFTEGKGVNVILDELGGPQMTLLGDVSATRGKL	240
PA1R_GCF_000496645.1|latest_PA1R_gp0285                EEREFLREQGADKVVVTDEQDLVLEVERFTEGKGVNVILDELGGPQMTLLGDVSATRGKL	240
                                                       ************************************************************

PAO1_Reference_GCF_000006765.1|latest_PA2491_mexS      VLYGCNGGDESAFPACAAFKKHLQFYRHCLMDFTGHPEMGLERNDESVSKALAHIEQLTR	300
LESB58_GCF_000026645.1|latest_PALES_28051              VLYGCNGGNESAFPACAAFKKHLQFYRHCLIDFTGHPEMGLERNDESVSKALAHIEQLTR	300
PAO1H2O_GCF_00071515.1|latest_OU9_02515                VLYGCNGGNESAFPACAAFKKHLQFYRHCLMDFTGHPEMGLERNDESVSKALAHIEQLTR	300
PAO1-VE2_GCF_000484495.2|latest_N296_2562              VLYGCNGGNESAFPACAAFKKHLQFYRHCLMDFTGHPEMGLERNDESVSKALAHIEQLTR	300
PAO1-VE13_GCF_000484545.2|latest_N297_2562             VLYGCNGGNESAFPACAAFKKHLQFYRHCLMDFTGHPEMGLERNDESVSKALAHIEQLTR	300
UCBPP-PA14_GCF_000014625.1|latest_PA14_32420           VLYGCNGGNESAFPACAAFKKHLQFYRHCLMDFTGHPEMGLERNDESVSKALAHIEQLTR	300
C7447m_GCF_000468935.2|latest_M802_2559                VLYGCNGGNESAFPACAAFKKHLQFYRHCLMDFTGHPEMGLERNDESVSKALAHIEQLTR	300
PA1_GCF_000496605.2|latest_PA1S_RS13175                VLYGCNGGNESAFPACAAFKKHLQFYRHCLMDFTGHPEMGLERNDESVSKALAHIEQLTR	300
PA1R_GCF_000496645.1|latest_PA1R_gp0285                VLYGCNGGNESAFPACAAFKKHLQFYRHCLMDFTGHPEMGLERNDESVSKALAHIEQLTR	300
                                                       ********:*********************:*****************************

PAO1_Reference_GCF_000006765.1|latest_PA2491_mexS      DRLLKPVVDRVFEFDQVVEAHRYMETCPKRGRVVIHVAD	339
LESB58_GCF_000026645.1|latest_PALES_28051              DRLLKPVVDRVFEFDQVVEAHRYMETCPKRGRVVIHVAD	339
PAO1H2O_GCF_00071515.1|latest_OU9_02515                DRLLKPVVDRVFEFDQVVEAHRYMETCPKRGRVVIHVA-	338
PAO1-VE2_GCF_000484495.2|latest_N296_2562              DRLLKPVVDRVFEFDQVVEAHRYMETCPKRGRVVIHVAD	339
PAO1-VE13_GCF_000484545.2|latest_N297_2562             DRLLKPVVDRVFEFDQVVEAHRYMETCPKRGRVVIHVAD	339
UCBPP-PA14_GCF_000014625.1|latest_PA14_32420           DRLLKPVVDRVFEFDQVVEAHRYMETCPKRGRVVIHVAD	339
C7447m_GCF_000468935.2|latest_M802_2559                DRLLKPVVDRVFEFDQVVEAHRYMETCPKRGRVVIHVAD	339
PA1_GCF_000496605.2|latest_PA1S_RS13175                DRLLKPVVDRVFEFDQVVEAHRYMETCPKRGRVVIHVAD	339
PA1R_GCF_000496645.1|latest_PA1R_gp0285                DRLLKPVVDRVFEFDQVVEAHRYMETCPKRGRVVIHVAD	339
                                                       **************************************
```

**Supplemental Figure S3: The XY and XXY forms of *mexT* in *P. aeruginosa* PAO1.** An 8 bp insertion causes a frame-shift in *mexT* within some *P. aeruginosa* PAO1 isolates, including the reference strain (PAO1-UW). This was masked by a mis-assigned upstream start codon in the sequences database (Winsor *et al.*, 2016). Removal of the 8 bp repeat in the isogenic mutant PA∆, restores the full length HTH domain, the correct reading frame and produces a functional protein. Protein domains were identified through sequence homology searches on InterPro (Pfam).

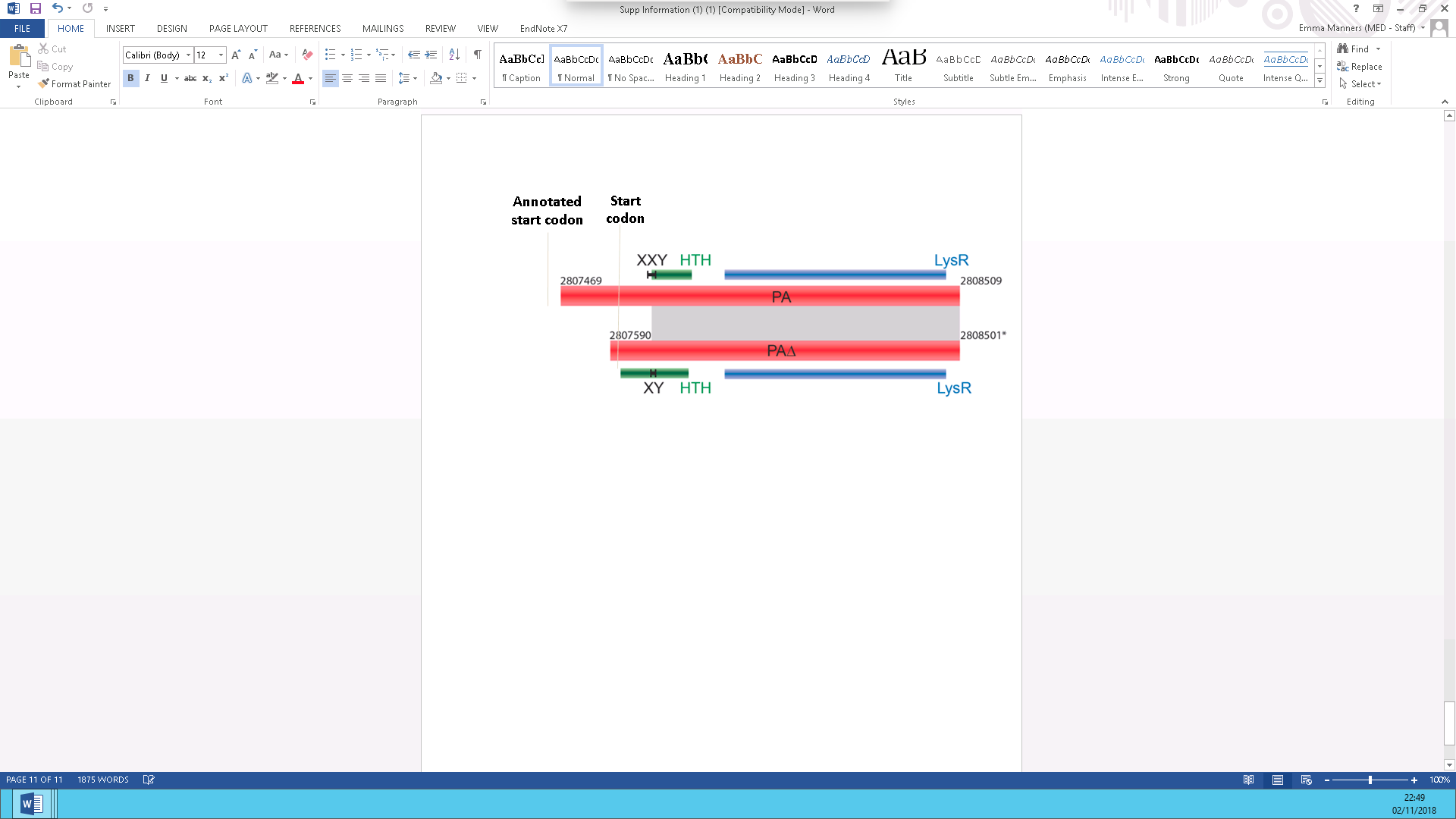

**Supplemental Figure S4: Growth of *P. aeruginosa* parent, mutant and revertant strains in media with and without chloramphenicol (8µg/ml).** A. Bacterial growth over time measured by the optical density at 600 nm. Two independent experiments (with five replicates) were performed; the bold line shows the average growth with error bars showing the 95% confidence intervals B. Box plots showing the area under the growth curves.

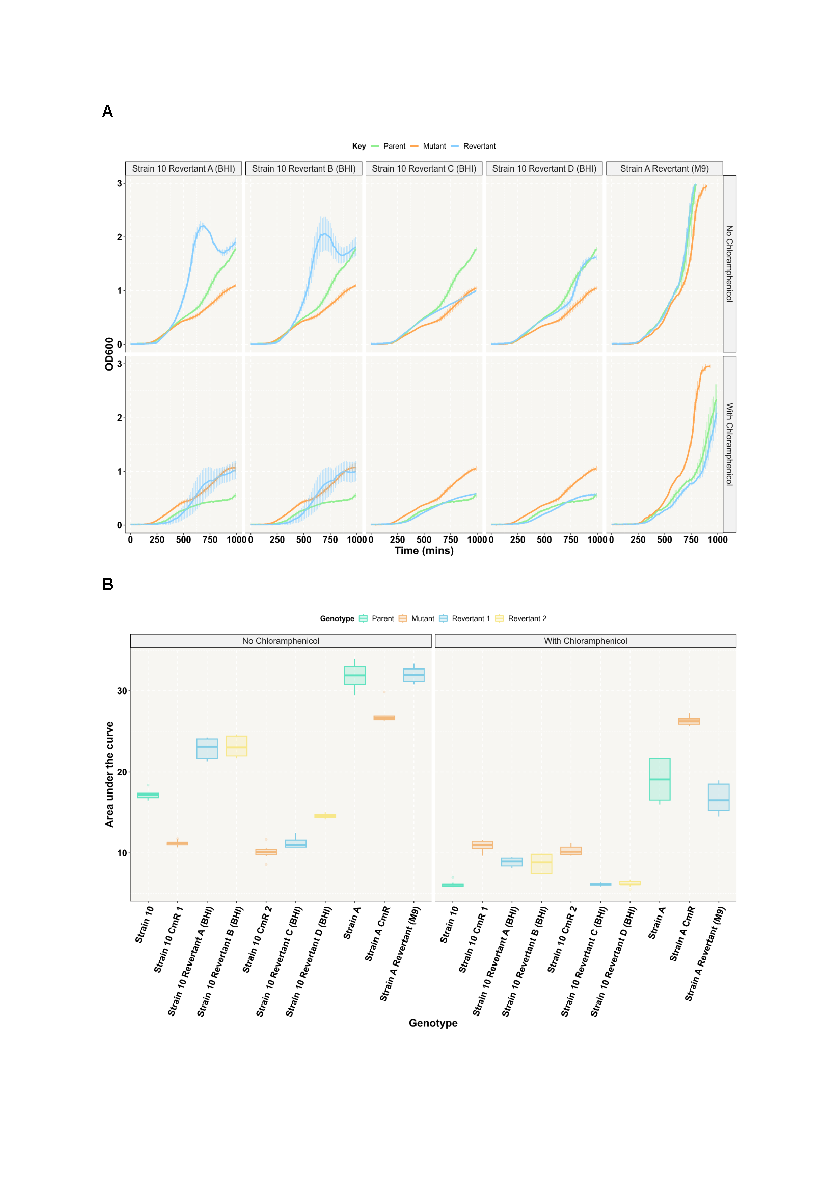

**Supplemental Figure S5:** Heatmap of the the data from dataset S5 focussed on the *mex* operon. The heatmap was formed from the dataset readout using R libraries to aggregate the data according to SNPEff variant type and provide total counts.

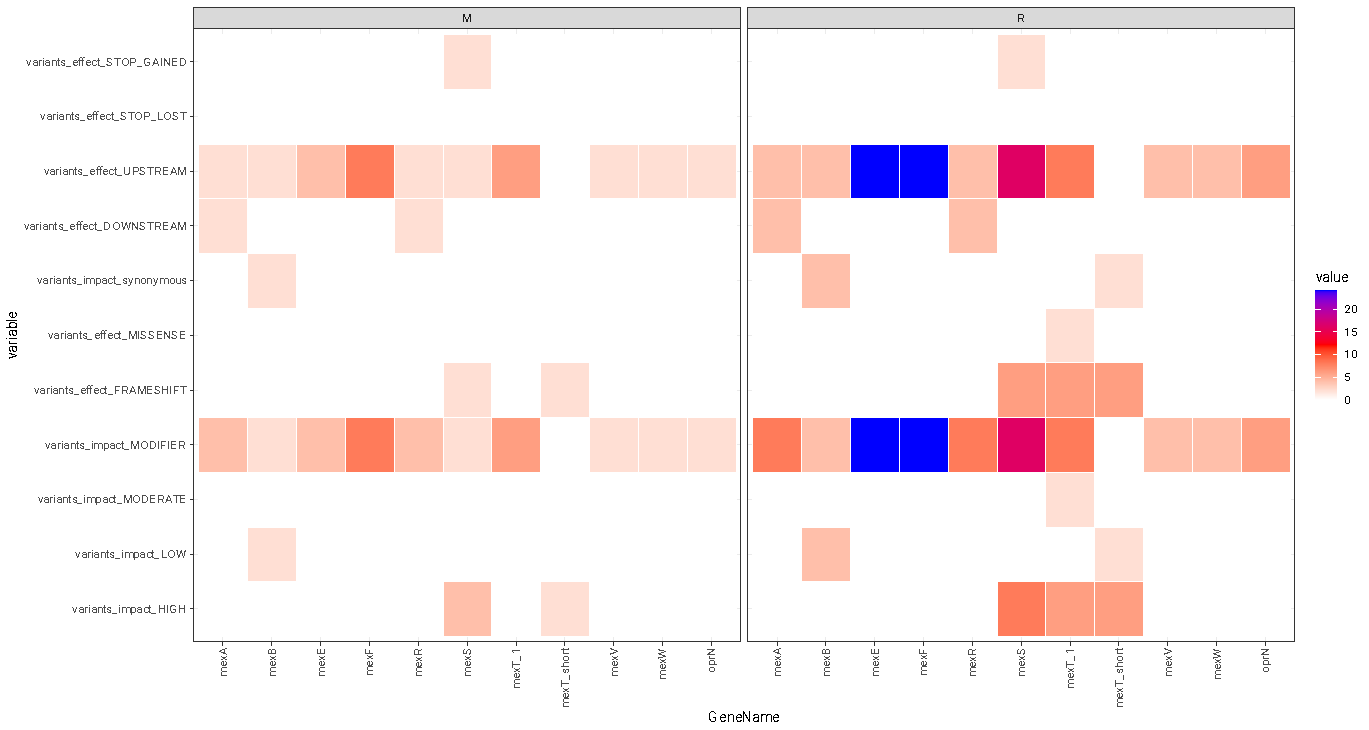
